# A growth-maintenance tradeoff determines nutrient-limited growth in phytoplankton

**DOI:** 10.64898/2026.06.01.729340

**Authors:** Ravi Ranjan, Alexey Ryabov, Kimberly Halsey, Helmut Hillebrand, Mridul K. Thomas, Bernd Blasius

**Author notes:** **Corresponding author email:** (Ravi Ranjan).

## Abstract

Phytoplankton encounter a range of light and nutrient conditions in nature and must adjust their internal carbon and nitrogen allocations to grow across different resource environments. Current phytoplankton carbon budget models treat respiration simply as a carbon loss. In reality, respiration is a critical cellular process that produces energy for nutrient uptake and cellular maintenance. Drawing on empirical evidence, we developed an eco-physiological model that incorporates a more realistic role of respiration. In our model, photosynthetic carbon is partitioned into: (i) the Pentose Phosphate Pathway (PPP) for assimilation and (ii) respiration for energy production that is then used in nutrient uptake. Stored nitrogen is partitioned between three pools: cellular structure, photosynthesis and nutrient uptake. Using an optimality-based approach, we identify strategies that maximize either exponential growth rate or competitive ability. We find that optimal internal allocations follow a growth-maintenance tradeoff, favoring population growth through carbon acquisition in nitrogen-replete conditions and population maintenance through nitrogen acquisition in nitrogen-limited conditions. The optimal allocations match empirically observed shifts in carbon partitioning at different dilution rates. Our model also generates an interactive growth response surface with an asymmetry, where light is the dominant limiting factor at low light intensities and co-limitation by light and nitrogen only occurs at high light levels. Furthermore, the model recovers the widely accepted Droop function for growth vs nitrogen quota and predicts a hyperbolic decline in growth vs energy quotas. Through a simple growth-maintenance tradeoff, our model provides a mechanistic foundation for predicting phytoplankton productivity in biogeochemical models.

## Introduction

‘To spend on one side, nature is forced to economize on the other side.’ This principle, famously cited by Darwin (Darwin 1861), has deep conceptual roots - tracing back to Aristotle (Aristotle 350 BC) and formalized in Goethe and Geoffroy’s *Loi de balancement* (law of balance) (Geoffroy Saint-Hilaire 1818). While originally invoked to describe morphological evolution, this economy-of-nature principle is equally critical in phytoplankton physiology. These microorganisms, responsible for half of Earth’s primary productivity, must constantly navigate this ‘law of balance’ by partitioning limiting internal resources such as carbon (*C*) and nitrogen (*N*) between the competing demands of growth and maintenance. This internal optimization is further complicated by a dynamic external environment, where nitrogen concentrations can vary over orders of magnitude and light needed for carbon assimilation vary within seconds to between years. Anthropogenic change is currently exacerbating nitrogen gradients (Jickells et al. 2017) and expanding oligotrophic zones (Leonelli et al. 2022), forcing phytoplankton to grow under increasingly variable conditions of nitrogen supply and fluctuating light. The varying external availability requires flexibility in the phytoplankton cell to optimally allocate the limited internal resources.

Current mathematical models of the phytoplankton carbon budget typically address this question through a rigid carbon balance, proposing that growth equals the difference between carbon uptake through photosynthesis and loss through respiration and excretion (Zonneveld et al. 1997; Pahlow 2005; Pahlow and Oschlies 2009). In this view, excretion is typically assumed to be small, and respiration is treated primarily as a fixed ‘tax’ proportional to maintenance. However, this framing of the carbon budget, with respiration solely being a cost, creates a biological paradox. It implies that growth would be maximized when respiration approaches zero. But a cell that does not respire is a dead cell. Respiration is more than just a carbon-loss process; it is the metabolic engine that catabolizes carbon to reclaim harvested light energy that has already been invested into carbon fixation. Thus, respiration effectively produces energy in the form of ATP, which is used by the cell for nutrient uptake, cellular maintenance and biosynthesis. Thus, even though respiration releases carbon back into the atmosphere, it is an indispensable *investment* of carbon required to acquire and process resources.

In addition to respiration, there is another major pathway in the carbon budget: the Pentose Phosphate Pathway (PPP) (Halsey et al. 2013, 2014; Halsey and Jones 2015). Through a series of chemostat experiments, Halsey and colleagues showed that the carbon acquired via photosynthesis (photosynthate from now onwards) is partitioned between respiration and PPP. Respiration catabolizes carbon to generate ATP needed for nutrient uptake, carbon concentration and maintenance activities. In contrast, the PPP catabolizes carbon to produce reductant that is essential in biosynthesis of lipids and nucleic acids during growth. In the Halsey experiments, the proportion of photosynthate allocated to the two pathways varied based on the dilution rates. When dilution rates were high and the population was forced to grow fast, phytoplankton species allocated less photosynthate to respiration and more photosynthate to PPP (Halsey et al. 2013). Halsey and colleagues hypothesized that faster growth rates led to more demand for reduced forms of carbons such as lipids and nucleic acids for cell division, therefore the cell allocated more photosynthate to PPP to produce the necessary reductant (Halsey and Jones 2015). Meanwhile, at lower growth rates, the cell was hypothesized to need more energy for maintenance (Geider and Osborne 1989; Kliphuis et al. 2012), favoring greater allocation of photosynthate toward respiration. These findings hint at a growth-maintenance tradeoff at play in the cellular carbon budget in phytoplankton.

Nitrogen is the primary constituent of proteins and nucleic acids in a phytoplankton cell. Since enzymes are proteins that modulate reaction rates, cellular nitrogen determines metabolic rates in phytoplankton. Nitrogen is taken up from the external environment through receptors, transported into the cell against an active concentration gradient using energy and allocated to metabolic pathways generating pigments for photosynthesis and enzymes for nutrient uptake. Furthermore, photosynthate partitioning into respiration vs PPP in the Halsey experiments was determined by nitrogen-limited growth rates. These results suggest that nitrogen partitioning is concomitant to carbon partitioning in phytoplankton, which judiciously partition both in response to the external environment. We hypothesized that when external nitrogen concentrations are low, more nitrogen is allocated to nitrogen uptake receptors to increase intracellular nitrogen and maintain the population. In contrast, nitrogen-replete conditions lead to allocation of nitrogen towards carbon uptake through photosynthesis for growth. Thus, both nitrogen and carbon cell budgets might have a concurrent growth-maintenance tradeoff at play.

Inspired by these experiments, we built a mathematical model to examine how optimal allocation of carbon and nitrogen determines the growth of phytoplankton across different nutrient and light environments. Our framework allows flexible allocation of both carbon and nitrogen to either growth-related processes (e.g., carbon intake and assimilation) or maintenance-related processes (e.g., maximizing nutrient uptake). We ask: what is the optimal allocation of carbon and nitrogen needed to maximize growth and how does it change with the external nitrogen environment? We also examine the optimal allocation to maximize competitive ability in a chemostat and analyze how the optimal allocations change with the dilution rate. Finally, we compare our model predictions regarding the optimal allocation at different dilution rates with the experimental photosynthate allocation patterns observed in the Halsey experiments.

We take an optimality-based approach to analyze our model (Litchman and Klausmeier 2008; Smith et al. 2011; Klausmeier et al. 2020). Optimality-based models typically define a measure of fitness, such as exponential growth rate, and identify the optimal trait value(s) that maximize fitness given biological tradeoffs. Such models have a long history in plankton ecology. Previous models have focused on a variety of ecological quantities in phytoplankton and examined a range of tradeoffs, including competition for light and nutrients (Litchman and Klausmeier 2001), using iron for light harvesting vs N assimilation (Armstrong 1999) and biosynthesis versus photosynthesis (Pahlow and Oschlies 2009). For more comprehensive reviews of optimality-based approaches in phytoplankton, see Geider et al. (2009) and Smith et al. (2011). However, unlike previous optimality-based models, our model explicitly incorporates the dynamic partitioning of photosynthate among competing metabolic pathways, allowing us to mechanistically connect environmental forcing to intracellular carbon allocation and growth. Thus, our model goes a step further from previous optimality models and links cellular biochemistry to population biology.

We analyze the model under two complementary conditions: (i) exponential growth limited by light and nutrients, and (ii) a light-saturated but nutrient-limited chemostat at equilibrium. Under the exponential growth condition, the phytoplankton growth occurs at a fixed light and nutrient level with no feedback between the population density and external nutrients. A culture experiences these conditions in a chemostat in the initial stage when a small inoculum is introduced and nutrient levels are at their input concentrations. At equilibrium, the growth rate in a chemostat is determined by the dilution rate through feedback between the external nutrient concentration and population density. We show that a growth-maintenance tradeoff mediates phytoplankton responses to external light and nitrogen availability under both conditions. We also derive a prediction for a phytoplankton growth surface over a light-nitrogen gradient. Under chemostat conditions, we demonstrate that the growth-maintenance tradeoff accounts for the patterns observed in the Halsey experiment, specifically the shifting allocation of photosynthate between respiration and PPP.

### Model description

We focus on the budgets of two important elements in a phytoplankton cell: carbon (*C*) and nitrogen (*N*). These budgets track the uptake and allocation of these elements and are linked through energy (*E*). Both carbon and nitrogen are taken up and allocated to competing metabolic processes. We analyze the model to determine the optimal allocation that either maximizes exponential growth or competitive ability.

A schematic of the model is shown in Fig. 1. Table 1 lists all the symbols used in the model along with their definitions and units. All analyses were done using the EcoEvo package (Klausmeier 2020) in Wolfram Mathematica 14.2.1 (Wolfram Research, Inc. 2024). In the next sections, we elaborate on the model and develop equations for each of the three state variables: assimilated carbon (*C*), nitrogen (*N*) and energy (*E*).

**Table 1:** Units and definitions of parameters and state variables.

| Symbol | Units | Detailed expression | Description |
| --- | --- | --- | --- |
| $C$ | $\text{mmol m}^{-3}$ | | Phytoplankton carbon concentration |
| $N$ | $\text{mmol m}^{-3}$ | | Phytoplankton nitrogen concentration |
| $E$ | $\text{mmol m}^{-3}$ | | Phytoplankton energy (ATP) concentration |
| $Q_N$ | $\text{mol N (mol C)}^{-1}$ | $N/C$ | Nitrogen quota |
| $Q_E$ | $\text{mol E (mol C)}^{-1}$ | $E/C$ | Energy quota |
| $N_{\text{str}}$ | $\text{mmol m}^{-3}$ | | Nitrogen allocated to structure |
| $N_{\text{nut}}$ | $\text{mmol m}^{-3}$ | | Nitrogen allocated to uptake |
| $N_{\text{photo}}$ | $\text{mmol m}^{-3}$ | | Nitrogen allocated to photosynthesis |
| $Q_{\text{str}}$ | $\text{mol N (mol C)}^{-1}$ | $N_{\text{str}}/C$ | Nitrogen quota allocated to structure |
| $Q_{\text{nut}}$ | $\text{mol N (mol C)}^{-1}$ | $N_{\text{nut}}/C$ | Nitrogen quota allocated to uptake |
| $Q_{\text{photo}}$ | $\text{mol N (mol C)}^{-1}$ | $N_{\text{photo}}/C$ | Nitrogen quota allocated to photosynthesis |
| $f_{\text{nut}}$ | — | $Q_{\text{nut}}/Q_N$ | Fraction of nitrogen quota allocated to nutrient uptake; optimization parameter |
| $f_{\text{photo}}$ | — | $1 - f_{\text{nut}} - \frac{Q_{\text{str}}}{Q_N}$ | Fraction of nitrogen quota allocated to photosynthesis |
| $S$ | $\text{mmol m}^{-3}$ | | External nitrogen concentration |
| $K_S$ | $\text{mmol m}^{-3}$ | | Half-saturation constant of nutrient uptake |
| $V_0$ | $\text{day}^{-1}$ | | Potential maximum nitrogen uptake rate |
| $V_{\text{max}}$ | $\text{mol N (mol C day)}^{-1}$ | $\frac{V_0}{\phi_N} f_{\text{nut}} Q_E$ | Maximum nitrogen uptake rate |
| $V$ | $\text{mol N (mol C day)}^{-1}$ | $V_{\text{max}}[f_{\text{nut}}, Q_E] \frac{S}{K_S + S}$ | Nitrogen uptake rate |
| $S_{\text{in}}$ | $\text{mmol m}^{-3}$ | | Nitrogen supply rate |
| $a$ | $\text{day}^{-1}$ | | Dilution rate |
| $I$ | $\mu\text{E m}^{-2}\text{day}^{-1}$ | | Light intensity |
| $K_I$ | $\mu\text{E m}^{-2}\text{day}^{-1}$ | | Half-saturation constant of photosynthesis |
| $P_0$ | $\text{mol C (mol C day)}^{-1}$ | | Potential photosynthesis rate |
| $P$ | $\text{mol C (mol C day)}^{-1}$ | $P_0 f_{\text{photo}} \frac{I}{K_I + I}$ | Photosynthesis rate |
| $f_{\text{PPP}}$ | — | | Fraction of carbon allocated to PPP |
| $\omega$ | $\text{mol C (mol C day)}^{-1}$ | $f_{\text{PPP}} P$ | Carbon allocated to PPP |
| $\rho$ | $\text{mol C (mol C day)}^{-1}$ | $(1 - f_{\text{PPP}}) P$ | Carbon allocated to respiration |
| $\phi_N$ | $\text{mol ATP (mol N)}^{-1}$ | | mol ATP used per mol of N taken up |
| $\phi_\rho$ | $\text{mol ATP (mol N)}^{-1}$ | | mol ATP generated per mol of C respired |
| $e_\omega$ | — | | Efficiency of carbon assimilation of the PPP |
| $\mu$ | $\text{d}^{-1}$ | $e_\omega \omega$ | Exponential growth rate |
| $l_C$ | $\text{d}^{-1}$ | | Carbon loss rate |
| $l_N$ | $\text{d}^{-1}$ | | Nitrogen loss rate |
| $l_E$ | $\text{d}^{-1}$ | | Energy loss rate |

**Figure 1:**
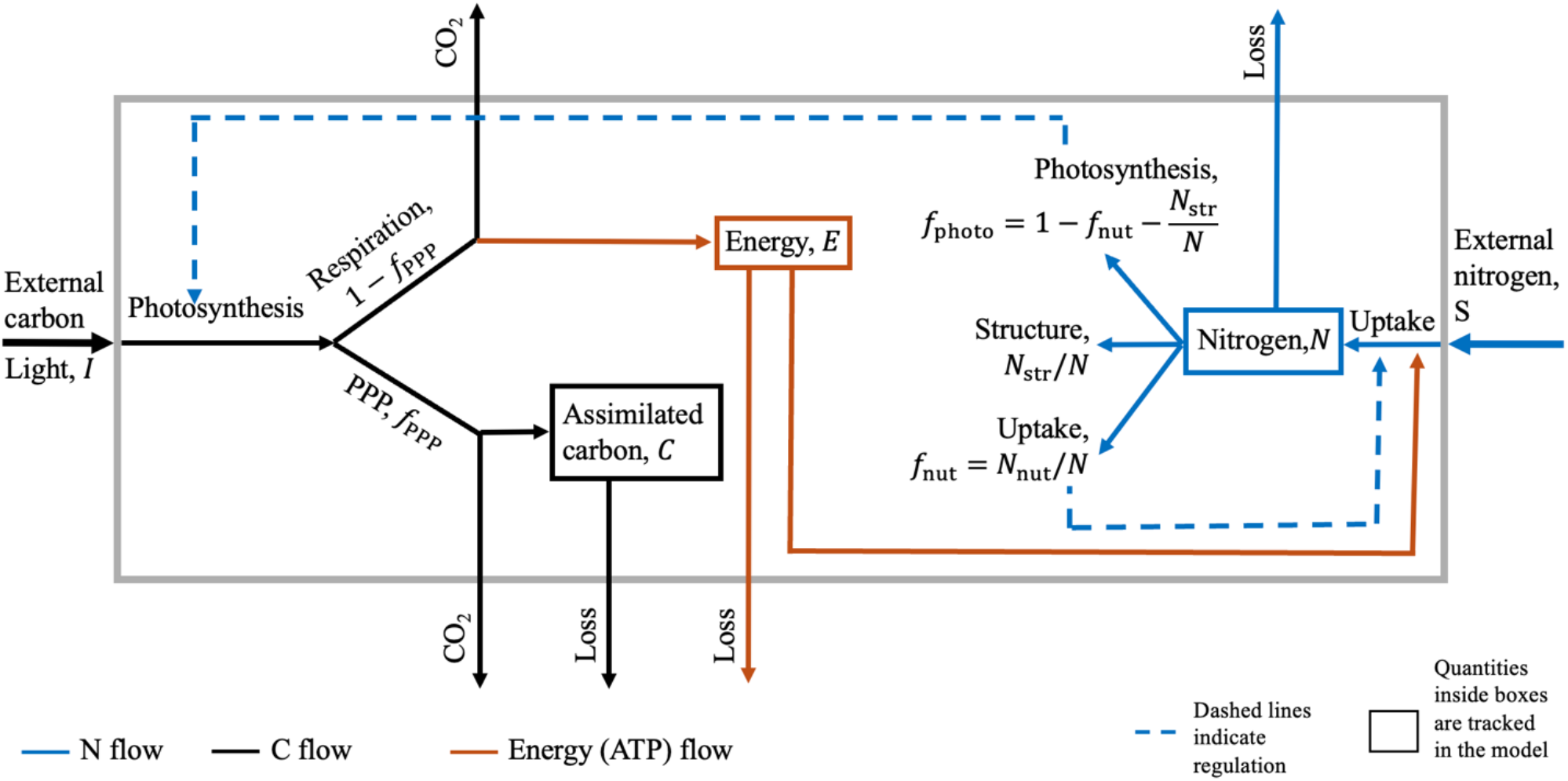
A schematic for the model denoting the nitrogen and carbon budgets and their interaction. External carbon (black arrows) is taken up via photosynthesis and allocated to two pathways: Pentose Phosphate Pathway (PPP; denoted as f_PPP_) and respiration (1 − f_PPP_). Respiration oxidizes the carbon into CO_2_ which exits the cell, and produces energy (E, brown arrows). PPP results in some carbon loss through oxidization to CO_2_ and the rest is assimilated carbon (C). External nitrogen (S, blue arrows) is taken up and stored in the cell. The stored nitrogen (N, blue arrows) is allocated three-ways: a fixed amount to structure (N_str_), a fraction to uptake (f_nut_ = N_nut_/N) and the remaining fraction to photosynthesis (f_photo_ = 1 − f_nut_ − N_str_/N). Nitrogen uptake consumes energy and is regulated by the nitrogen allocation to uptake. Nitrogen allocation to photosynthesis regulates photosynthesis rates. f_nut_ and f_PPP_ are the allocations that are optimized to maximize exponential growth or competitive ability.

#### Nitrogen dynamics

Nitrogen is taken up from the external environment and stored inside the cell. The flow of nitrogen is shown via blue arrows in Fig. 1. We assume that nitrogen concentration is proportional to enzyme concentration. Therefore, intracellular nitrogen concentrations regulate reaction rates in the model. The total intracellular nitrogen (*N*) is partitioned three-ways: photosynthesis (*N*_photo_), nutrient uptake (*N*_nut_) and structural nitrogen (*N*_str_),

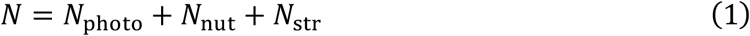

In terms of nitrogen cell quota (N:C ratio of the cell, *Q*_*N*_) this reads as *Q*_*N*_ = *Q*_photo_ + *Q*_nut_ + *Q*_str_, which can be rewritten as:

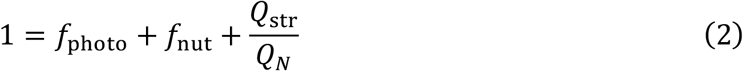

where *f*_photo_ = *Q*_photo_/*Q*_*N*_ and *f*_nut_ = *Q*_nut_/*Q*_*N*_. Since cellular structure is essential to the integrity of the cell, we assume that the nitrogen quota allocated to structure (*Q*_str_) is fixed (Pahlow and Oschlies 2013). The fraction of nitrogen quota allocated to nutrient uptake, *f*_nut_ is a species-specific allocation parameter that we later optimize for exponential growth or competitive ability. Since the total nitrogen quota *Q*_*N*_ is dynamic and depends on the external nitrogen concentration, the fraction of nitrogen allocated to photosynthesis *f*_photo_ is also dynamic and can be expressed as a function of *f*_nut_:

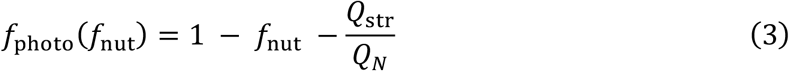

This has some important consequences. Most notably, to have a positive photosynthesis rate (*f*_photo_ > 0), the cell needs a minimum nitrogen quota, 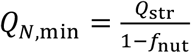. Secondly, when nitrogen quota becomes very large (*Q*_*N*_ → ∞), the fraction of nitrogen allocated to structure (*Q*_str_/*Q*_*N*_) approaches zero. This means that the fraction of nitrogen allocated to photosynthesis (*f*_photo_) asymptotically approaches the fraction of nitrogen quota remaining after allocation to uptake

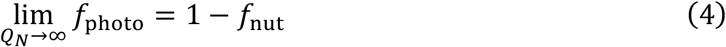

To derive the dynamic equation for nitrogen (*N*), we assume that the nitrogen concentration in the cell increases through uptake, *V*(*f*_nut_)*C*, and decreases due to a loss with fixed rate (*l*_*N*_).

Thus, the specific change rate of intracellular nitrogen is

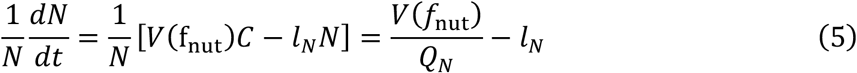

Here, we assume that nitrogen is taken up at the maximum uptake rate, *V*(*f*_nut_) = *V*_max_(*f*_nut_). We relax this assumption later. Nitrogen uptake is an energy-intensive process, therefore the maximum uptake rate *V*_max_ is determined by both the fraction of nitrogen allocation to the uptake machinery *f*_nut_ and the available energy quota *Q*_*E*_ = *E*/*C* (see below). We assume a multiplicative linear dependence on both factors

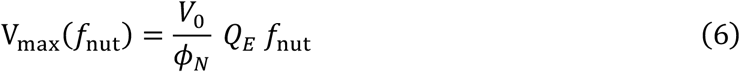

where *V*_0_ is the potential maximum uptake rate and *ϕ*_*N*_ is the energetic cost of transporting and assimilating one mol of nitrogen.

#### Carbon dynamics

External carbon enters the phytoplankton cell via photosynthesis (*P*). Unless indicated otherwise, we assume that the population is growing under saturated light conditions. The photosynthesis rate, *P*, is determined by the nitrogen allocation to the photosynthetic machinery (*f*_photo_), which in turn depends on the allocation to nutrient uptake, *f*_nut_, as defined in Eq 1. This leads to the following expression

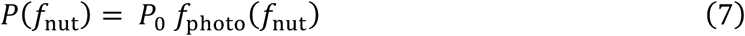

where *P*_0_ represents the potential maximum photosynthesis rate.

Following the empirical results of Halsey et al. (2013, 2014), the photosynthate (carbon produced by photosynthesis) is consumed by two primary pathways: respiration (*ρ*) and PPP (*ω*). We assume that the photosynthate is at steady state, meaning that the total photosynthate consumption equals production (*P* = *ρ* + *ω*). We denote the fraction of photosynthate allocated to the PPP as *f*_PPP_, with the remaining fraction (1 − *f*_PPP_), directed towards respiration (see Fig.1). Consequently, the reaction rate of respiration is given by:

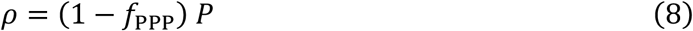

and the reaction rate of PPP is:

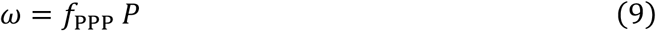

Similar to *f*_nut_, we treat *f*_PPP_ as a species trait to be optimized.

Both respiration and PPP are complex pathways involving several enzymes and intermediates some of which are also exchanged between the two pathways. However, to keep the model simple, we assume that they operate independently. Specifically, all carbon entering a pathway remains sequestered within the pathway in until fully oxidized, with no intermediate exchange between the two. Under these assumptions, the cell loses all carbon entering the respiration pathway in the form of CO_2_. In contrast, the PPP has an efficiency *e*_*ω*_ of assimilation: for every mole of carbon entering the pathway, 1 − *e*_*ω*_ moles are lost as CO_2_, while the remainder is assimilated.

The specific rate of carbon assimilation into biomass is thus

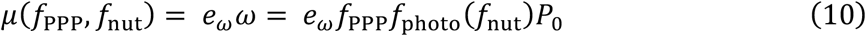

Let *C* denote the cellular pool of assimilated carbon, the net carbon balance is then expressed as

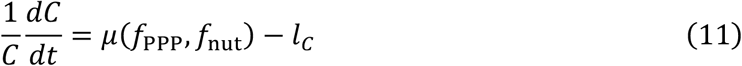

where *l*_*C*_ represents a specific carbon loss rate (e.g., via exudation).

#### Energy dynamics

Energy in the form of ATP is produced through the respiration pathway with a yield of *ϕ*_*ρ*_ mol ATP per mol C. Conversely, energy (ATP) is consumed during nutrient uptake since nutrients must be transported against a concentration gradient via active transport with an energetic cost of *ϕ*_*N*_ mol ATP per mol N. As with carbon and nitrogen, there is a fixed loss rate *l*_*E*_, yielding the energy balance

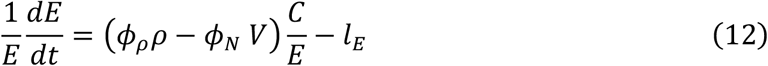

where *ρ* = (1 − *f*_PPP_) *P*_0_ *f*_photo_(*f*_nut_) is the reaction rate of respiration and *V* = *V*(*f*_nut_) the nitrogen uptake rate (see above).

#### Model in terms of intensive variables

We rewrite the model in terms of assimilated carbon, *C*, and quotas of nitrogen (*Q*_*N*_ = *N*/*C*) and energy (*Q*_*E*_ = *E*/*C*) quotas (see Section 1, Supplementary Information (SI)). As intensive variables, these quotas are easier to work with and are also ecologically relevant as a measure of nitrogen and energy storage. We further simplify the model by assuming that all loss rates are equal (*l*_*C*_ = *l*_*N*_ = *l*_*E*_). Under these assumptions the model can be written as:

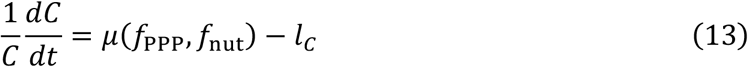

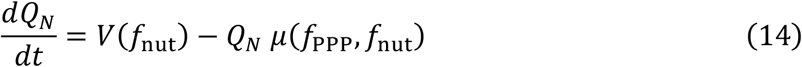

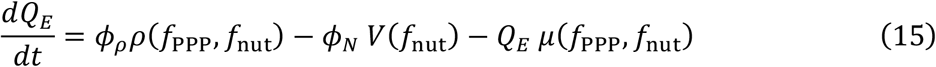

Our model builds upon the framework of Pahlow and colleagues (Pahlow 2005; Pahlow and Oschlies 2009, 2013) by incorporating carbon partitioning concurrently with nitrogen partitioning and explicitly tracking energy dynamics in the cell. Despite its simplicity, the model captures key trade-offs in carbon, nitrogen and energy allocation. Ecologically, the equations offer a transparent interpretation: net carbon dynamics reflect the balance between assimilation and losses; the nitrogen quota increases through uptake but is diluted by carbon assimilation; and the energy quota increases via respiration while being consumed by both nutrient uptake and carbon assimilation. All metabolic rates are expressed in terms of the two allocation parameters: carbon allocation to PPP (*f*_PPP_) and nitrogen allocation to uptake (*f*_nut_).

The allocations of carbon and nitrogen combine to generate either a population growth or a population maintenance strategy. Increasing the nitrogen allocated to photosynthesis (*f*_photo_(*f*_nut_)) and the photosynthate allocated to PPP (*f*_PPP_) together increases assimilated carbon and thus is an allocation towards population growth. Increasing nitrogen allocation to uptake (*f*_nut_) and carbon allocation to respiration (1 − *f*_PPP_) increases nutrient uptake rate and contributes to maintaining the population in low-nutrient conditions.

We then ask: **what are the optimal** 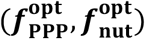 **values under different growth regimes such as exponential growth or a chemostat?** Furthermore, how do these optimal 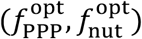 values change as external nitrogen or light conditions change?

### Model analysis

#### Balanced exponential growth

##### Calculation of balanced growth steady quota

We first examine the model under balanced exponential growth, the most studied physiological state in laboratory experiments with acclimated phytoplankton cultures. Under balanced exponential growth, all extensive state variables (carbon (*C*), nitrogen (*N*) and energy (*E*)) grow at the same constant rate. Consequently, the intensive state variables (quotas *Q*_*N*_ and *Q*_*E*_) remain constant over time:

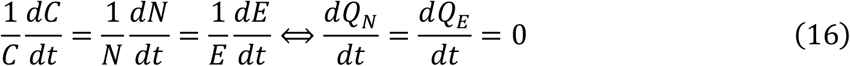

Assuming a saturated nutrient environment (*V* = *V*_max_), we solve these equations to obtain the non-trivial balanced growth (steady) quota values 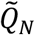 and 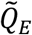 (see Section 1, SI for full expressions). We then use 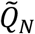 and 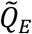 to express the exponential growth rate in terms of our two allocation traits: *f*_PPP_, and *f*_nut_. The trait *f*_PPP_ determines the fraction of photosynthate allocated to PPP and consequently to growth (Fig. 1). The trait *f*_nut_ determines the fraction of nitrogen quota allocated to nutrient uptake. Maximizing nutrient uptake is essential in times of low ambient nutrient levels, so a *f*_nut_ allocation contributes towards population maintenance. In this section, we calculate the optimal allocation levels 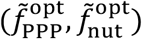 that maximize exponential growth. The parameter values used for generating the figures are listed in Table S1.

First, we examine how changing carbon and nitrogen allocations affect the steady nitrogen and energy quotas (Fig. 2). These results illustrate how the interaction between nutrient and carbon allocation regulates exponential growth. Increasing *f*_PPP_ (moving to the right on the x-axis in Figs. 2a, b) increases allocation towards growth but reduces respiration and energy production, as only the fraction 1 − *f*_PPP_ of photosynthate is directed towards respiration (Eq. 8). Consequently, the energy quota, 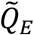, declines (Fig. 2b). This reduction in available energy suppresses nitrogen uptake, leading to a concurrent decrease in the nitrogen quota, 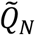 (Fig. 2a). These declines in steady quota span several orders of magnitude (note the logarithmic scaling in Fig. 2).

**Figure 2:**
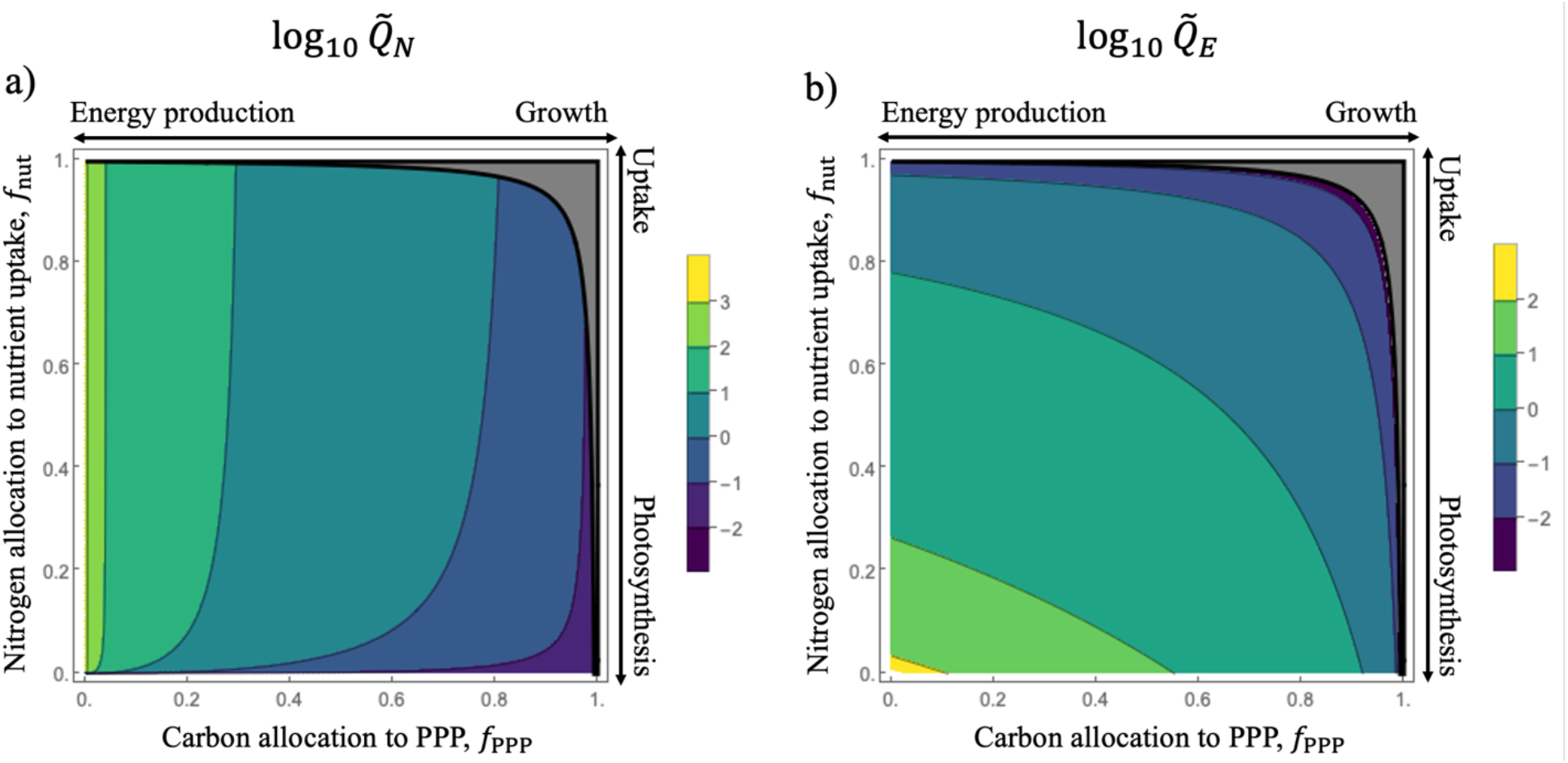
Influence of allocation parameters on steady nitrogen and energy quotas during balanced exponential growth. a) Log_10_-transformed steady nitrogen quota, 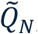, and b) energy quota, 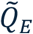, derived from Eqns. S11-12 in Section 1, SI. Values are shown as a function of the allocation parameters f_PPP_ and f_nut_. The grey region delineated by thick black lines represents a region of infeasible allocation parameters where growth is not possible because 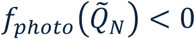, leading to negative 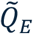.

Conversely, increasing *f*_nut_ (moving upwards on the y-axis of Figs. 2a, b) increases allocation towards nutrient acquisition, which increases energy consumption. Furthermore, higher *f*_nut_ reduces the nitrogen fraction allocated to photosynthesis *f*_photo_ (see Eq. 1), thus reducing carbon fixation. This reduction in carbon intake further limits respiration and energy production. The combination of high energy demand and low production result in a depletion of the energy storage at high *f*_nut_ values (Fig. 2b). While an initial rise in *f*_nut_ increases the nitrogen quota 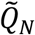 through enhanced uptake, the concurrently falling energy levels as *f*_nut_ continues to rise eventually causes 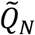 to saturate (Fig. 2b).

The top right corner in Figs. 2a, b (grey region with black boundary lines) corresponds to a strategy of simultaneous high carbon allocation to growth (*f*_PPP_) and high nitrogen allocation to uptake (*f*_nut_). However, balanced exponential growth is biologically infeasible in this region due to the exhaustion of nitrogen available for photosynthesis. For feasible exponential growth, the nitrogen fraction allocated to photosynthesis at steady state must be positive, such that 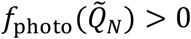. Thus, increasing *f*_nut_ directly reduces the available photosynthesis fraction 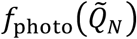. In contrast, increasing *f*_PPP_ has an indirect effect on *f*_photo_: by diverting carbon from respiration, it lowers energy production, which in turn suppresses nitrogen uptake and reduces nitrogen quota, 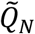. Consequently, high values of *f*_PPP_ lead to a lower nitrogen quota, ultimately lowering the fraction allocated for photosynthesis. When the nitrogen pool is insufficient to support photosynthesis, energy production fails 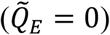 and the population goes extinct. In the model, the infeasible region (grey region in Figs. 2a, b) corresponds to the trivial solution of eq. 16 where 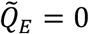.

##### Optimal allocation for exponential growth

Next, we evaluate the steady-state exponential growth rate 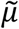 as a function of *f*_PPP_ and *f*_nut_ (Fig.3a). As established in Fig. 2b, growth is not possible in regions where 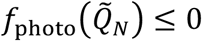. Within the feasible parameter space (non-grey region in Fig. 3a), the growth rate exhibits a unimodal response to both *f*_PPP_ and *f*_nut_. To explore this further, in Figs. 3b and 3c we plot cross-sections of the growth surface from Fig. 3a. In Fig. 3b, increasing *f*_PPP_ initially enhances the carbon flow through PPP, thereby increasing 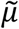. However, as discussed above, high *f*_PPP_ values indirectly deplete the nitrogen allocated to photosynthesis, eventually causing 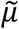 to decline. A similar trade-off is observed for *f*_nut_ (Fig. 3c). At low *f*_nut_ levels, nitrogen uptake is insufficient, leading to a low nitrogen quota, 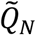. While initial increases in *f*_nut_ levels increase 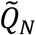 and the fraction available for photosynthesis 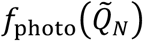, further increases in *f*_nut_ reduce *f*_photo_. Consequently, the growth rate reaches a maximum and subsequently declines.

**Figure 3:**
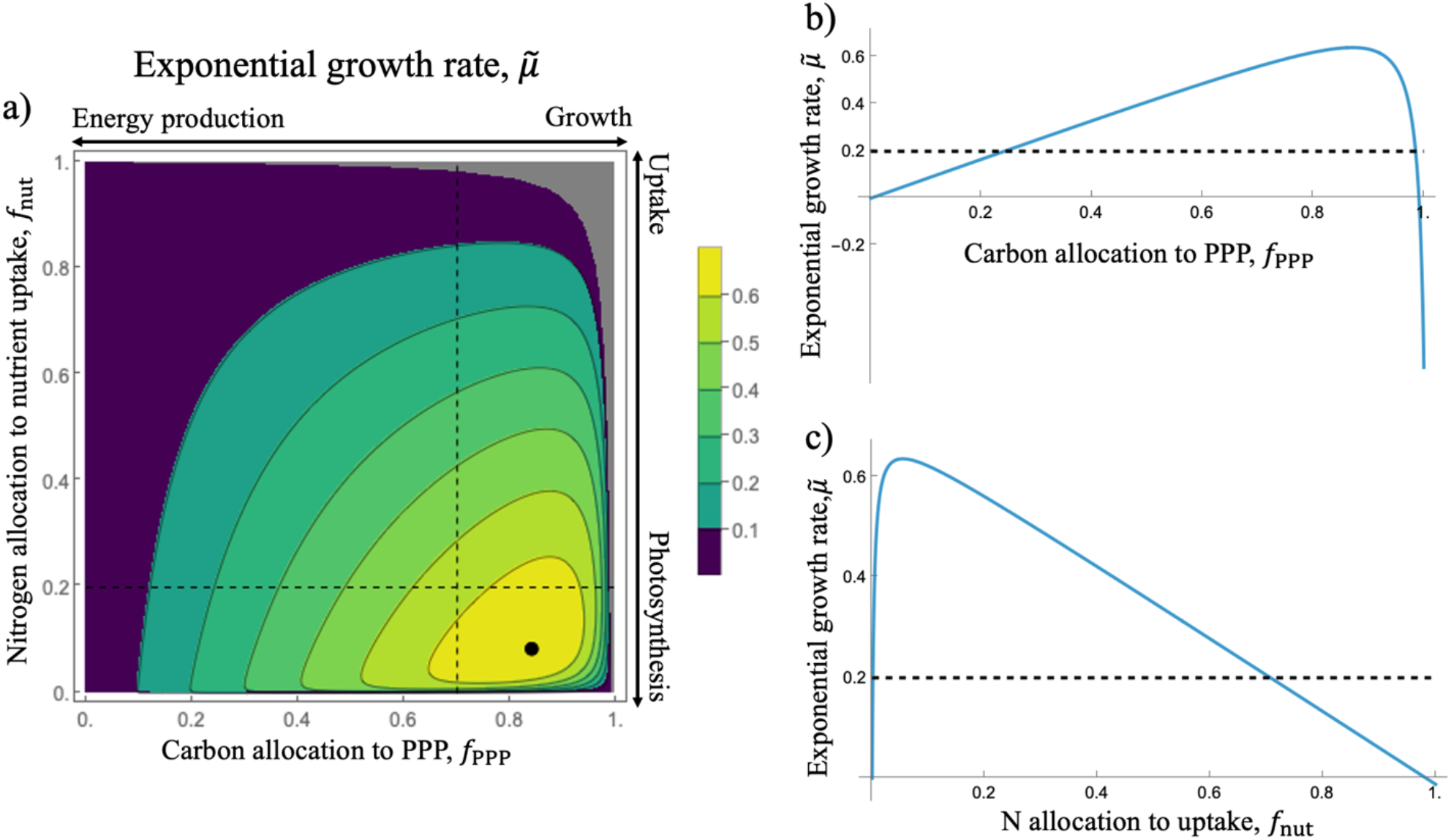
Dependence of the exponential growth rate, 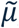, on the allocation parameters. a) Growth rate 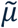 as a function of f_PPP_ and f_nut_. The grey region represents the parameter space where growth is infeasible (cf. Fig. 2). The black dot identifies the optimum allocation strategy 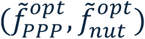 – the combination of allocation parameters that maximizes exponential growth rate. Dashed lines represent the cross sections shown in b) and c). b) Growth rate 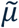 as a function of carbon allocation to PPP (f_PPP_) at a fixed nitrogen allocation (f_nut_ = 0.2). c) Growth rate 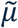 as a function of the nitrogen allocation to uptake (f_nut_) at a fixed carbon allocation (f_PPP_ = 0.7). The horizontal dashed lines in b) and c) denote the specific loss rate, l_C_ = 0.2.

The black dot in Fig. 3a indicates the optimal allocation strategy 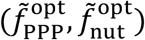 – the specific combination of *f*_PPP_ and *f*_nut_ that maximizes the balanced exponential growth rate. We determine this optimal strategy by calculating the local maximum of 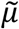 with respect to the steady quotas 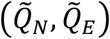 (details in Section 1, SI). When resources are non-limiting, optimal growth is achieved through a strategy of low *f*_nut_ and high *f*_PPP_, reflecting a prioritized investment in growth over maintenance. In the following section, we examine how these optimal allocations shift under light- or nutrient-limited conditions.

##### The impact of nutrient and light limitation on optimal allocations

Next, we relax the assumptions of light and nutrient saturation by calculating the optimal growth rate as a function of external light intensity and nutrient concentration. To do this, we replace the potential carbon uptake rate *P*_0_ in eq. 7 with a light-dependent uptake such that

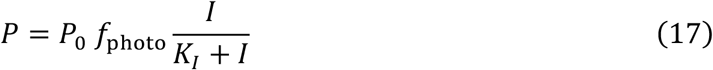

where *I* denotes the light intensity and *K*_*I*_ is the half-saturation constant. Similarly, we replace the potential nitrogen uptake rate, *V*_0_, with a nutrient-dependent uptake:

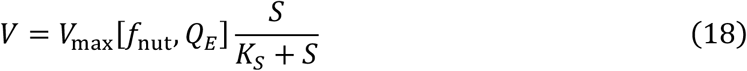

where *S* represents the external nitrogen concentration and *K*_*S*_ is its corresponding half-saturation constant.

The steady-state quotas, 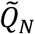 and 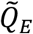, can be expressed as functions of *f*_PPP_, *f*_nut_ and the ratio of carbon and nitrogen uptake rates *P*/*V* (Section 2, SI). Substituting these quotas into the growth rate expression and maximizing it with respect to (*f*_PPP_, *f*_nut_) yields the optimal allocations 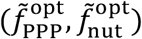 and the corresponding optimal growth rate 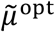 as functions of *P*/*V*. Consequently, the optimal allocations and the optimal growth rate can also be written as a function of the light intensity *I* and the nitrogen concentration *S*. To maintain balanced growth, the cell adjusts its optimal allocations to account for mismatches in the carbon vs nitrogen uptake rates (Fig. S1; see Section 2, SI for a thorough analysis). This internal modulation of the optimal allocations drives the growth rate in the light and nitrogen limited case described below.

In Fig. 4a, we illustrate the optimal growth rate as a function of light intensity, *I*, and nutrient concentration, *S*. The growth rate increases with both light intensity and nutrient concentration in an interactive way. This interaction is further illustrated in Fig. 4c, which shows that growth saturates with increasing light for any given nutrient concentration. Notably, with higher nutrient concentrations the growth-light curve reaches saturation at a higher growth rate and has a higher half-saturation constant for light intensity (Fig. 4e). We observe a reciprocal pattern in Fig. 4d: growth saturates with nutrient concentrations like a Monod curve. Thereby, the highest achieved growth rate and the half-saturation constant both increase with light intensity (Fig. 4f).

**Figure 4:**
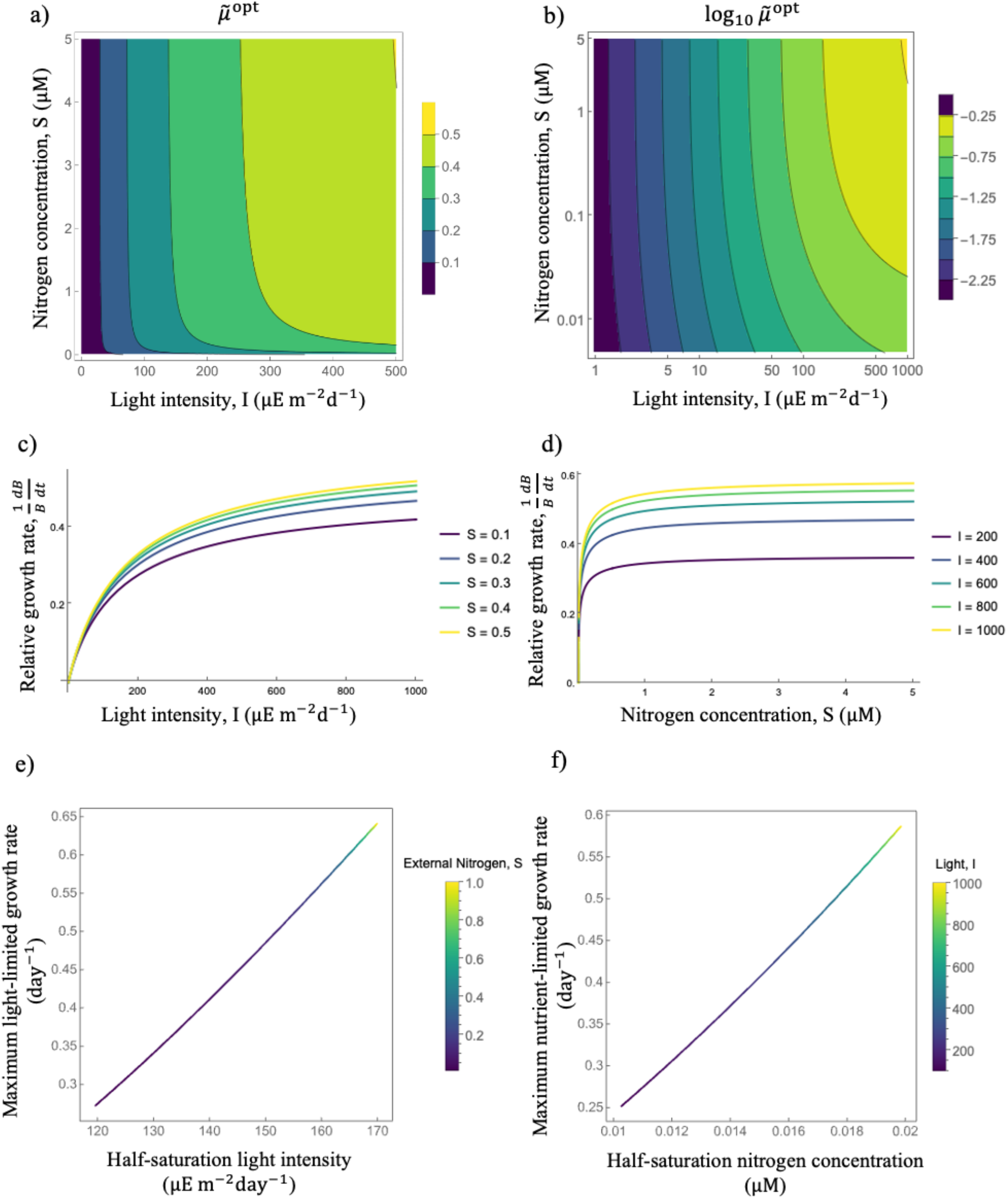
Interactive effects of light intensity and nitrogen concentration on the optimal growth rate. a) Optimal growth rate, μ, shown in color as a function of light intensity I and nitrogen concentration S. Warmer colors indicate higher growth rates. b) Same as a) using a logarithmic representation, showing the common logarithm (log_10_) of the growth rate as a function of light intensity I and nitrogen concentration S on a log-log scale. c) Relative growth rate as a function of light at several nitrogen concentrations. Increasing S increases both the maximum growth rate and the light half-saturation constant K_J_. d) Relative growth rate as a function of nitrogen at several light intensities I. Similarly, higher light intensity leads to an increase in the maximum growth rate and the nitrogen half-saturation constant K_K_. e) Increasing nitrogen concentration (S, shown by changing colors of the curve) increases both maximum growth rates (y-axis) and half-saturation constants (x-axis) of the light-limited growth curve in c. f) Increasing light intensity (I, shown by changing colors of the curve) increases both maximum growth rates (y-axis) and half-saturation constants (x-axis) of the nitrogen-limited growth curve in d.

The log-log representation in Fig. 4b reveals the nature of the limitation regimes. At low light intensities, the growth contours are almost vertical, indicating that the system is nutrient-saturated but primarily light-limited. As light intensity increases, the system transitions into a regime where both nutrients and light are co-limiting. This asymmetric limitation pattern in our eco-physiological model contrasts with standard models that use a product of two single-driver Monod functions, which typically assume co-limitation at all light levels.

Our model also predicts that nutrient-limited growth reaches saturation more rapidly than light-limited growth (Figs. 4a-d). This occurs because external nitrogen influences growth only indirectly through the internal allocation of carbon and nitrogen. Therefore, even a high *K*_*K*_ for nitrogen uptake does not significantly elevate the half-saturation constant for N-limited growth. In contrast, light exerts both a direct impact on carbon fixation through photosynthesis and an indirect impact through internal allocation of carbon and nitrogen, causing the limit-limited growth curve to track the underlying light-limited carbon uptake curve more closely.

#### Chemostat dynamics

##### Optimal allocation for competitive equilibrium

Thus far, we have examined the model under exponential growth without feedback between the population density and resource availability. We now consider a chemostat environment under saturated light, where the phytoplankton growth rate is determined by the dilution rate, *D*, set by the experimenter. To achieve equilibrium, the population reduces the ambient nitrogen concentration to *S*^∗^, the minimum level at which growth rate matches dilution (*μ* = *D*). *S*^∗^ measures the competitive ability for the nutrient: a species with a lower *S*^∗^ can deplete the nutrient to a level that excludes competitors with higher nutrient requirements. Since growth rate is now determined externally by the dilution rate, we cannot optimize for growth rate in a chemostat. Instead, we now optimize for *S*^∗^. Specifically, we ask: what is the optimal allocation of carbon to PPP (*f*_PPP_) and nitrogen to nutrient uptake (*f*_nut_) that minimizes *S*^∗^, thereby defining the most competitive strategy for a given dilution rate?

Ambient nitrogen, *S*, is supplied with a supply rate, *S*_in_, and the dilution rate is set at *D*. Assuming phytoplankton nitrogen uptake follows a Monod-type response, the nitrogen dynamics are described as:

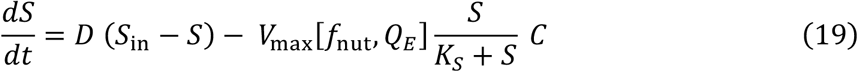

The remaining model equations stay the same as in the previous sections, but we set all loss rates equal to the dilution rate (*l*_*C*_ = *l*_*N*_ = *l*_*E*_ = *D*) since dilution is the primary driver of loss in a chemostat. We calculate the equilibrium values 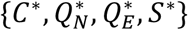 by setting the time derivatives to zero:

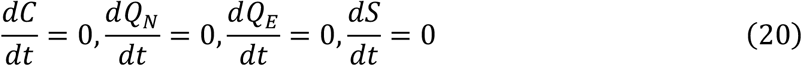

The full analytic expressions for 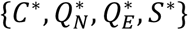 are provided in Section 4, SI. We begin our analysis by examining how the internal quotas of nitrogen and energy shift in response to changes in the allocation parameters.

In Fig. 5a, we plot the logarithm-transformed (base 10) nitrogen quota at equilibrium, 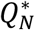, as a function of the two allocation parameters *f*_PPP_ and *f*_nut_. The grey region, delineated by dashed brown lines, represents an infeasible parameter space where the nitrogen quota 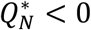. This occurs because dilution exceeds the maximum attainable growth. At low *f*_PPP_ levels (the left part of the infeasible region), insufficient carbon is allocated to biomass synthesis via the PPP to offset the losses from dilution. Conversely, at high *f*_nut_ values (the top part of the infeasible region), excessive nitrogen investment in uptake machinery limits the nitrogen available for photosynthesis. In both cases, the resulting growth rate falls below the dilution rate, making a steady-state population unsustainable.

**Figure 5:**
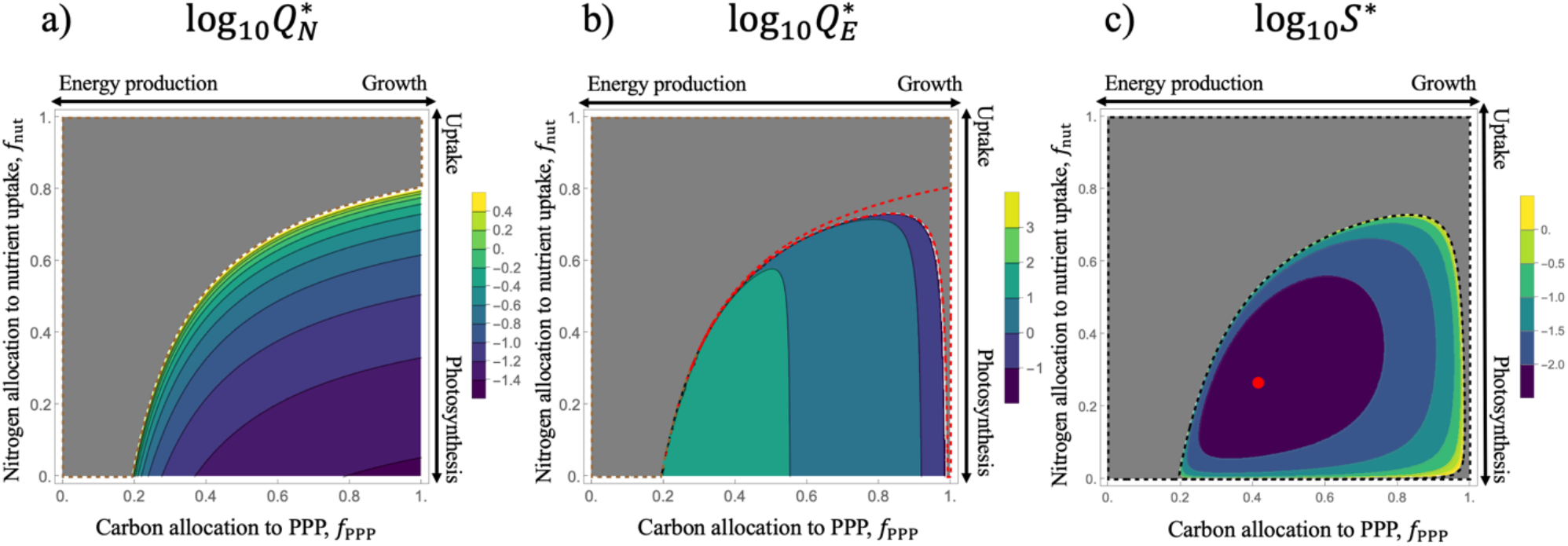
Effects of allocation parameters on quotas and minimal nitrogen requirements in a chemostat. Steady-state values of a) nitrogen quota 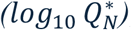, b) energy quota 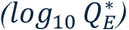, and c) minimum nitrogen requirement (log_10_ S^∗^) as a function of carbon and nutrient allocation at D = 0.2. In all panels, grey regions represent infeasible allocation values. The grey regions are delineated in a) by brown dashed lines 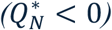 and in b) by red dashed lines 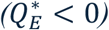. The infeasible region defined by S^∗^ (S^∗^ < 0) is the overall infeasible region and encompasses the infeasible regions from a) and b). It is delineated in c) by black dashed lines. The red dot in c) indicates the optimal allocation of carbon and nitrogen for maximizing competitive ability.

As *f*_PPP_ increases beyond the threshold needed to match growth to the dilution rate (moving to the right in Fig. 5a), the carbon flux through the PPP goes up while respiration declines. This reduction in respiration leads to lower energy production, which in turn suppresses nitrogen uptake and results in a lower steady-state nitrogen quota, 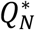 (cf. Fig.1). Within the feasible region 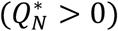, increasing the nitrogen allocation to uptake, *f*_nut_, (moving upward along the y-axis) initially enhances nitrogen uptake and storage. The positive relationship between 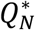 and *f*_nut_ persists until the nitrogen remaining for photosynthesis becomes insufficient to sustain the population against dilution (dashed brown curve in Fig. 5a).

Next, we plot the steady-state energy quota 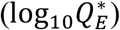 against the allocation parameters *f*_PPP_ and *f*_nut_ (Fig. 5b). Colored areas indicate feasible equilibria 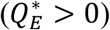. These are not restricted only by the previously identified nitrogen-limited boundary (brown lines), but also by a second infeasible region (red boundary) where 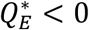. Within this red-bounded region, high *f*_PPP_ suppresses respiration and thus energy production, while high *f*_nut_ elevates the energy demand for nutrient uptake. In both scenarios, energy consumption exceeds production, leading to 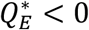 and precluding a steady state.

Within the feasible colored region in Fig. 5b, 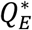 decreases as allocation shifts toward higher *f*_PPP_ (reducing carbon available for respiration) or higher *f*_nut_ (increasing energy expenditure for uptake). While *Q*_*E*_ and *Q*_*N*_ exhibit similar sensitivity to allocation during both exponential growth and at equilibrium, the chemostat environment imposes more stringent constraints. As can be seen by comparing Figs. 2 and 5, exponential growth is feasible for most of the parameter space except at very high values of *f*_PPP_ and *f*_nut_. In contrast, the chemostat equilibrium requires the population growth rate to match the dilution rate *D*. This additional constraint renders large sections of the allocation space, which are viable during exponential growth, infeasible at equilibrium.

Finally, we examine the equilibrium resource concentration (log_10_ *S*^∗^) as a function of the allocation parameters *f*_PPP_ and *f*_nut_ (Fig. 5c). The black dashed lines represent the most restrictive feasibility boundary (*S*^∗^ < 0) and the infeasible region within them subsumes the previous two infeasible regions (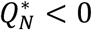 and 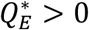). We observe a non-monotonic relationship between *f*_PPP_ and *S*^∗^. At low *f*_PPP_ (left boundary of feasible region), carbon allocation towards growth is minimal. To satisfy the chemostat requirement (*μ* = *D*), this low growth efficiency must be compensated by a higher amount of carbon uptake through photosynthesis. The low *f*_PPP_ allocation leads to increased energy production and nitrogen storage 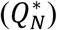, which enhances *f*_photo_ and the resulting photosynthesis rate. Due to the high 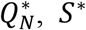 remains high at low *f*_PPP_values. Conversely, at the right boundary (high *f*_PPP_), suppressed respiration rates limit energy availability for nutrient uptake, again resulting in elevated *S*^∗^. Therefore, *S*^∗^ is minimized at intermediate values of *f*_PPP_, where the balance between growth investment and energy production optimizes competitive ability.

A similar non-monotonic pattern is observed along the *f*_nut_ axis (Fig. 5c). At low values of *f*_nut_ (bottom of the feasible region), the nutrient uptake rate is low, leading to a higher equilibrium resource concentration *S*^∗^ to sustain growth. Conversely, at high *f*_nut_ values (top of the feasible region), the excessive allocation to uptake reduces the nitrogen available for photosynthesis. But the high allocation to uptake also increases the nitrogen quota 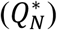 which bolsters *f*_photo_ and in turn elevates *S*^∗^. Therefore, *S*^∗^ is minimized at intermediate values of *f*_nut_, reflecting the trade-off between maximizing nutrient acquisition and maintaining the photosynthetic machinery.

The optimal competitive strategy 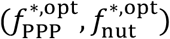, defined at the allocation parameters that minimize *S*^∗^, is marked by the red point in Fig. 5c. At this coordinate the species achieves its highest competitive ability; according to *R*^∗^ theory, no other strategy can invade or outcompete this optimum at the given dilution rate. Because the chemostat equilibrium requires *μ* = *D*, the optimal allocation is intrinsically dependent on the dilution rate. To investigate this dependency, we calculate the optimal strategy by solving for the critical points where ∇*S*^∗^(*f*_PPP_, *f*_nut_) = 0. We verify that the critical point is a local minimum by ensuring the Hessian matrix of *S*^∗^ is positive definite (i.e., a positive determinant and the top left element of the Hessian is positive). The complete analytical expressions for these optimal allocations are provided in Section 4, SI.

Contrasting how the equilibrium resource concentration (*S*^∗^, Fig. 5c) and the exponential growth rate (*μ*, Fig. 3a) depend on the allocations is instructive. Since *D* = 0.2 in Fig. 5, the boundary of feasible region for an equilibrium in Fig. 5c is the same as the contour line for *μ* = 0.2 in Fig.3a. The optimal allocation strategy to maximize exponential growth increases growth through a high *f*_PPP_ and a low *f*_nut_ so that *f*_photo_ is high. However, in a chemostat, the optimal strategy is to have a low allocation to *f*_PPP_ and a slightly higher allocation to *f*_nut_ so that *S*^∗^ is minimized. Thus, while the *μ*-optimal strategy might grow faster during periods of replete nutrients, it will be out-competed by a *S*^∗^-optimal strategy at equilibrium.

##### The role of dilution rate

In Fig. 6a, we plot the optimal competitive allocation parameters as a function of the chemostat dilution rate. To maintain equilibrium as the dilution rate increases, the population must elevate its growth rate. Mechanistically, this can be achieved in two ways, either increasing total carbon fixation through photosynthesis or increasing the fraction of photosynthetic carbon going towards growth (PPP). We find that as dilution rates increase, both occur. Thereby the model predicts a strategic shift in nitrogen allocation: optimal *f*_nut_ decreases, allowing for a corresponding increase in *f*_photo_, thus elevating the photosynthesis rate. Concurrently, the fraction of carbon going towards growth also increases due to an increased optimal *f*_PPP_. Thus, the cell adjusts both carbon and nitrogen allocation to meet the increased growth demand imposed by the higher turnover rate.

**Figure 6:**
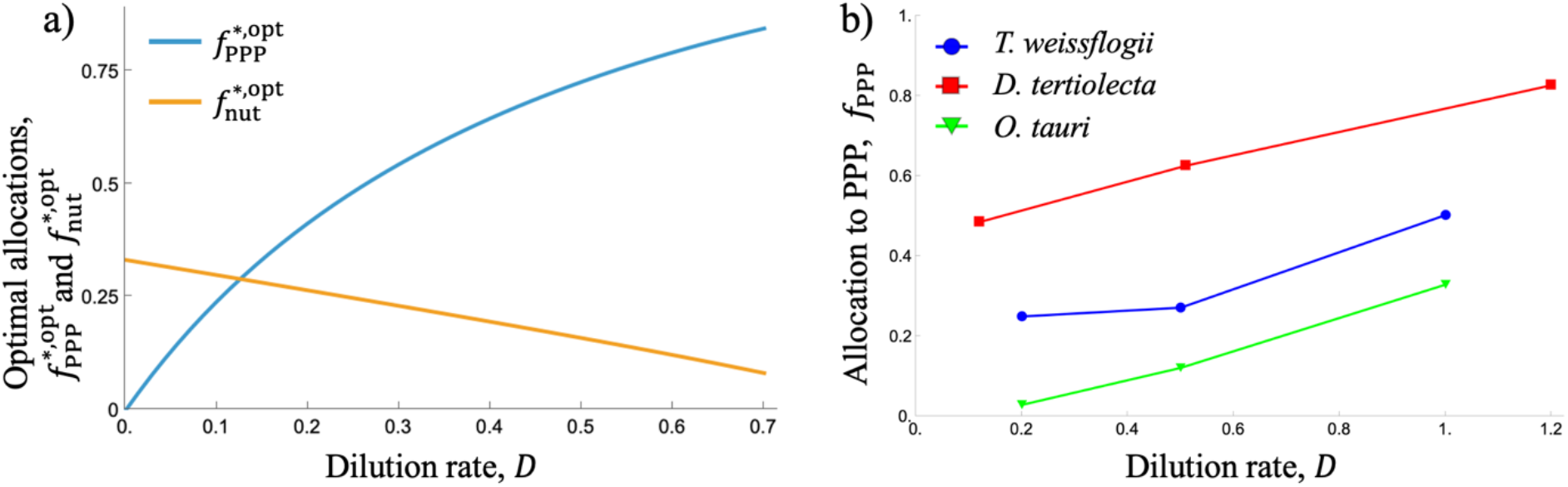
Influence of dilution rate on optimal allocation in a chemostat and empirical validation. a) Optimal allocation of carbon to PPP 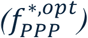 and nitrogen to uptake 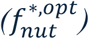 at equilibrium as a function of the dilution rate, D. b) Comparison with experimental data (adapted from Halsey et al. 2013, 2014) showing that the fraction of carbon directed towards PPP (light blue region) increases with dilution rate for three evolutionarily distant phytoplankton species: Thalassiosira weissflogii (blue), Dunaliella tertiolecta (red) and Ostreococcus tauri (green).

To validate our model predictions, we compared our results with chemostat experiments conducted by Halsey et al. (2013, 2014). In these studies, three evolutionarily distant phytoplankton species, the chlorophyte *Dunaliella tertiolecta*, the diatom *Thalassiosira weissflogii* (Halsey et al. 2013), and the picoeukaryotic prasinophyte *Ostreococcus tauri* (Halsey et al. 2014), were grown across three dilution rates. At every dilution rate the authors measured gross and net carbon production, defining their difference as transient carbon - the fraction of fixed carbon that exits the cell over the course of a single cell cycle. This transient pool was further partitioned into carbon flux directed toward the PPP and carbon lost through mitochondrial respiration. We extracted these carbon production data using the digitizing tool PlotDigitizer (plotdigitizer.com) from Fig. 1 in Halsey et al. (2013) and Fig. 2 in Halsey et al. (2014).

While the empirical carbon production data are normalized by absorbed light, our model scales production by total cellular carbon in the population. To reconcile these different scaling factors, we focus on the fractional allocation towards PPP, which is scale-invariant and thus facilitates a direct comparison. We calculated the fraction of transient carbon (defined as Gross Primary Production minus Net Primary Production; GPC – NPC) that is directed towards PPP. In Fig. 6b, we plot this fraction across the three experimental dilution rates (0.2, 0.5, 1.0 d ^−1^ for *T. weissflogii* and *O. tauri*; 0.12, 0.51, 1.2 d ^−1^ for *Dunaliella tertiolecta*). Consistent with our model’s predictions, the proportion of carbon directed toward the PPP increases with the dilution rate, while the proportion of carbon lost to respiration decreases. This trend also appears to hold for *Micromonas pusilla*, although the data for this species were less amenable to reliable extraction.

## Discussion

Intracellular carbon and nitrogen in phytoplankton are governed by a trade-off between population growth vs population maintenance. In resource-replete environments, fitness is maximized through investment in cell division and population growth. Conversely, resource-limited environments favor investment in the uptake machinery to sustain the population. The growth-maintenance tradeoff emerges from a combination of two underlying partitions: carbon towards respiration (maintenance) vs PPP (growth) and nitrogen towards uptake (maintenance) vs photosynthesis (growth). We formalized these partitions using two key allocation traits: the fraction of carbon allocated towards PPP (*f*_PPP_) and the fraction of nitrogen allocated towards nutrient uptake (*f*_nut_). We then examined the implications of this tradeoff under two growth regimes: exponential growth and a chemostat. In both regimes, we show that the stoichiometric coupling between these pathways renders “single-purpose” strategies, which maximize one function at the total expense of the other, physiologically infeasible. Instead, we derive optimal allocation strategies that maximize either the intrinsic rate of increase during exponential growth or competitive ability at equilibrium, with profound impacts for cellular energy and nitrogen quotas.

The exponential growth rate depends unimodally on both carbon allocation to PPP (*f*_PPP_) and nitrogen allocation to uptake (*f*_nut_), resulting in a growth optimum at intermediate allocation values. Increasing *f*_PPP_ initially increases growth by accelerating biomass synthesis, however excessive carbon allocation to PPP eventually becomes self-limiting, as it reduces carbon available for respiration and suppresses the energy supply required for nitrogen uptake. Similarly, increasing *f*_nut_ initially enhances growth through higher nitrogen storage, but further investment reduces the nitrogen allocation for photosynthesis. This reduction in the photosynthetic machinery ultimately constrains the carbon supply, leading to a decline in the overall growth rate.

The optimal allocation strategy shifts in response to light and nitrogen availability to maintain maximum growth (see Section 2, SI for a complete exploration). Under high light intensity but low nitrogen concentrations, the cell prioritizes nitrogen acquisition by co-regulating both traits to alleviate the nutrient limitation. Vice versa, when nitrogen is abundant but light is limiting, the system shifts toward maximizing both total carbon fixation through increased photosynthetic investment and the efficiency of biomass production through *f*_PPP_. Therefore, these internal allocation traits act as a coordinated regulatory mechanism that allows the cell to counteract stoichiometric mismatches in the external carbon and nitrogen supply.

By incorporating these optimal allocations, we evaluated the exponential growth rate as a function of light and nutrient availability. Notably, our model reveals an interesting asymmetry in the sensitivity to light versus nutrients. At low light intensity, growth is primarily light-limited and nutrient saturation occurs at exceptionally low nitrogen concentrations. Co-limitation of growth only emerges under high-light conditions. The more limiting impact of light is to be expected: while nitrogen availability influences growth mostly indirectly through shifts in optimal allocation, light impacts growth both directly through carbon fixation and indirectly by modulating the metabolic state and the resulting allocation strategies.

While the separate dependencies of growth on light and nutrients are well-understood, their possible interaction remains a subject of active debate. Our model provides a mechanistic, biologically grounded prediction of the growth surface, derived from first-principles allocation trade-offs. By capturing the interactive effects of light intensity and nutrient availability, this framework provides a robust foundation for biogeochemical models to calculate phytoplankton productivity across the diverse light and nutrient regimes encountered in global oceans.

Our model reduces to a Droop-like function for the relationship between growth rate and internal nitrogen levels (Section 3, SI). Previously, Pahlow and Oschlies (2013) demonstrated that the Droop model can be derived from an optimality-based approach focusing on carbon and nitrogen allocation. We extend this theoretical foundation by showing that the Droop functional form can be derived from a more complex metabolic architecture that incorporates explicit energy dynamics and the simultaneous optimization of both carbon and nitrogen. Our model also predicts a hyperbolic decline in growth as energy storage increases, with a critical minimum energy quota, 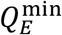, required for viable growth. In this context, elevated energy storage serves as a physiological indicator of high carbon allocation toward respiration, thus reducing the flux toward PPP and constraining the maximum achievable growth rate.

Under chemostat conditions, the predicted shifts in carbon and nitrogen allocation at different dilution rates reflect a fundamental growth-maintenance tradeoff. At low dilution rates, the optimal strategy prioritizes maintenance and nitrogen acquisition to sustain the population under severe limitation. This is achieved by diverting more carbon towards respiration (to fuel uptake) and more nitrogen toward the uptake machinery. As the dilution rate increases, the equilibrium growth requirement (*μ* = *D*) forces the population to grow quicker and shifts both carbon and nitrogen allocation towards biosynthetic pathways, increasing the fractions dedicated to the PPP and photosynthesis, respectively.

Our model shows that maximizing competitive ability in a chemostat requires a starkly different allocation strategy than maximizing exponential growth. The growth-optimal opportunist strategy has a high carbon allocation towards PPP and a low nitrogen allocation towards uptake. In contrast, a gleaner strategy with a low PPP allocation and increased uptake allocation is optimal for a chemostat. Thus, our model provides a physiological basis for the gleaner-opportunist tradeoff which has been widely studied in the context of species coexistence (Armstrong and McGehee 1980; Chesson 2000; Litchman and Klausmeier 2001; Tsakalakis et al. 2018). While the gleaner-opportunist tradeoff is typically thought of in terms of a single limiting resource like nitrogen (Grover 1990; Swanson et al. 2024), our model shows that it involves both carbon and nitrogen at the physiological level. The opportunist strategy is essentially a C-acquisition strategy that succeeds when nitrogen is replete. In contrast, the gleaner strategy is a N-acquisition strategy that succeeds when nitrogen is limiting. The cross-elemental nature of the tradeoff may also be the reason why previous efforts to find the tradeoffs focusing on just carbon acquisition traits failed (Swanson et al. 2024).

Our model predictions regarding carbon allocation at different dilution rates are consistent with the experimental observations of Halsey and Jones (2015). Their study demonstrated that as dilution rates increase, the allocation of transient carbon (carbon that enters and leaves the cell within a cell cycle) shifts significantly: investment in the PPP increases while the fraction diverted toward respiration decreases. These conserved patterns across evolutionarily distant phytoplankton species led Halsey et al. to hypothesize that the ATP:NADPH ratio of the cell acts as the primary metabolic driver, specifically that respiration provides the high ATP demand of maintenance at slow growth, whereas the PPP supplies the reductant required for rapid biosynthesis. While our model accounts for respiratory energy production, it purposefully omits explicit tracking of reductant (NADPH) production. Instead, by representing respiration as a dual process of carbon loss and energy production and the PPP as a carbon-accumulation flux, we demonstrate that a conceptually simpler growth-maintenance trade-off is sufficient to recover these empirical trends. This suggests that the observed allocation shifts may emerge from fundamental stoichiometric constraints rather than specific redox-ratio regulation alone.

Our framework incorporates several simplifying assumptions regarding the biochemistry of the phytoplankton cell. We represent respiration and the PPP as independent linear pathways whose reaction rates solely depend on the photosynthate concentration. We also assume that carbon molecules entering either of these pathways complete the entire pathway. In reality, both respiration and PPP are complex metabolic networks involving numerous intermediates and shared metabolites. As stated earlier, we also omit the specific role of reductants (e.g., NADPH) in these dynamics. However, the fundamental behavior of our model hinges on the carbon-loss and energy production resulting from respiration and carbon-accumulation resulting from PPP. Therefore, as long as respiration is the higher carbon-loss process and PPP is the higher carbon-accumulating process, the model results will still hold. While the addition of finer metabolic complexities would likely refine the quantitative sensitivity of the allocation trends, the predicted directionality of the growth-maintenance trade-off would remain unchanged.

Our findings suggest several promising avenues for future research. Incorporating more realism by exploring the consequences of unequal losses of carbon and nitrogen through exudation will be important in the context of global carbon and nitrogen cycles (Thornton 2014). While we assumed a constant chlorophyll-to-carbon ratio to focus on the allocation of photosynthetic carbon between competing pathways, it is well-established that phytoplankton dynamically adjust their pigment content in response to light and nitrogen environments (photoacclimation). A key next step will be to integrate these acclimation dynamics with our growth-maintenance trade-off model. Developing an optimality-based framework that simultaneously accounts for pigment regulation and carbon-nitrogen allocation will be key towards effective prediction of the biological carbon pump and its response to a changing ocean.

## Acknowledgements

No conflicts of interest. RR acknowledges support from the HIFMB Postdoc Program (HIPP) at HIFMB, a collaboration between the Alfred-Wegener-Institute, Helmholtz-Center for Polar and Marine Research and the Carl-von-Ossietzky University Oldenburg, initially funded by the Ministry of Science and Culture of Lower Saxony (MWK) and the Volkswagen Foundation through the ‘Niedersächsisches Vorab’ grant programme (ZN3285). AR was funded by the European Union under the Horizon Europe program (grant agreement no. 101081642, www.obama-next.eu).

## Data Accessibility Statement

No new data was generated during the current study. The data used in Fig. 6 is included in the accompanying Mathematica notebook (link in the Supplementary Information).

## Notes

### Competing Interest Statement

The authors have declared no competing interest.

### Summary of Updates

Removing couple of pages not relevant for the preprint.

