## Supplementary Information for "A growth-maintenance tradeoff determines nutrient-limited growth in phytoplankton"

**Supporting Information for:** A growth-maintenance tradeoff determines nutrient-limited growth in phytoplankton

**Authors:** Ravi Ranjan[1,2,3,4,*] (ORCID: 0000-0001-8644-9316), Alexey Ryabov[2,5] (ORCID: 0000-0002-1595-6940), Kimberly Halsey[6] (ORCID: 0000-0002-1407-0777), Helmut Hillebrand[1,2,3,5] (ORCID: 0000-0001-7449-1613), Mridul K. Thomas[7] (ORCID: 0000-0002-5089-5610) and Bernd Blasius[5] (ORCID: 0000-0002-6558-1462)

[1] Helmholtz Institute for Functional Marine Biodiversity at the University of Oldenburg, Oldenburg, Germany;

[2] Alfred Wegener Institute, Helmholtz Centre for Polar and Marine Research, Bremerhaven, Germany

[3]Hanse-Wissenschaftskolleg Institute for Advanced Study, Delmenhorst, Germany

[4] Department of Integrative Biology, University of Texas at Austin, Austin, Texas 78751 USA

[5] Institute for Chemistry and Biology of the Marine Environment, University of Oldenburg, Carl-von-Ossietzky-Straße 9-11, 26129 Oldenburg, Germany

[6] Department of Microbiology, Oregon State University, 2820 SW Campus Way, Corvallis, OR 97331, USA

[7]Department F.-A. Forel for Environmental and Aquatic Sciences and Institute for Environmental Sciences, University of Geneva, Geneva, Switzerland;

**Running head:** Phytoplankton carbon budget model

### Supplementary Information

Main text: Ranjan R, Ryabov A, Halsey K, Hillebrand H, Thomas MK, Blasius B. A growth-maintenance tradeoff determines nutrient-limited growth in phytoplankton. bioRxiv. https://doi.org/10.64898/2026.06.01.729340

Code: https://github.com/raviranjan545/Carbon_Budget_Model

#### 1. Model analysis during nitrogen-replete exponential growth

The model tracks three extensive variables (carbon ($C$), nitrogen ($N$) and energy ($E$)), whose dynamics are described by:

$$\frac{1}{C}\frac{dC}{dt} = \mu(f_{\mathrm{PPP}}, f_{\mathrm{nut}}) - l_C \tag{S1}$$

$$\frac{1}{N}\frac{dN}{dt} = \frac{V(f_{\mathrm{nut}})}{Q_N} - l_N \tag{S2}$$

$$\frac{1}{E}\frac{dE}{dt} = \left(\phi_\rho \rho - \phi_N\, V(f_{\mathrm{nut}})\right)\frac{C}{E} - l_E \tag{S3}$$

These equations are the same as eqns. 5, 11 and 12 in the main text. The model has two allocation parameters: nitrogen allocation to uptake machinery, $f_{\mathrm{nut}}$, and carbon allocation to the Pentose Phosphate Pathway (PPP), $f_{\mathrm{PPP}}$. The fraction of nitrogen allocated to photosynthesis ($f_{\mathrm{photo}}$) is:

$$f_{\mathrm{photo}}(f_{\mathrm{nut}}) = 1 - f_{\mathrm{nut}} - \frac{Q_{\mathrm{str}}}{Q_N} \tag{S4}$$

Here, $\mu(f_{\mathrm{PPP}}, f_{\mathrm{nut}})$ is the specific rate of carbon assimilation into biomass:

$$\mu(f_{\mathrm{PPP}}, f_{\mathrm{nut}}) = P_0\, e_\omega\, f_{\mathrm{PPP}}\, f_{\mathrm{photo}}(f_{\mathrm{nut}}) \tag{S5}$$

The rate of nitrogen uptake is $V(f_{\mathrm{nut}})$. We start with the assumption that the nitrogen uptake rate is the maximum uptake rate $\mathrm{V}_{\max}(f_{\mathrm{nut}})$, which depends on $f_{\mathrm{nut}}$ and $Q_E$:

$$V(f_{\mathrm{nut}}) = \mathrm{V}_{\max}(f_{\mathrm{nut}}) = \frac{V_0}{\phi_N}\, Q_E\, f_{\mathrm{nut}} \tag{S6}$$

We assume that the loss rates are equal for all three state variables: $l_C = l_N = l_E$. We rewrite the models in terms of the two intensive variables: nitrogen quota ($Q_N = N/C$) and the energy quota ($Q_E = E/C$).

$$\frac{dQ_N}{dt} = \frac{1}{B}\left(\frac{dN}{dt} - \frac{N}{B}\frac{dB}{dt}\right) = V(f_{\mathrm{nut}}) - Q_N\, \mu(f_{\mathrm{PPP}}, f_{\mathrm{nut}}) \tag{S7}$$

$$\frac{dQ_E}{dt} = \frac{1}{B}\left(\frac{dE}{dt} - \frac{E}{B}\frac{dB}{dt}\right) = \phi_\rho \rho(f_{\mathrm{PPP}}, f_{\mathrm{nut}}) - \phi_N\, V(f_{\mathrm{nut}}) - Q_E\, \mu(f_{\mathrm{PPP}}, f_{\mathrm{nut}}) \tag{S8}$$

During balanced exponential growth, all components of the population grow at the same rate. In the model, this means that the carbon ($C$), nitrogen ($N$) and energy ($E$) grow at the same rate.

$$\frac{1}{C}\frac{dC}{dt} = \frac{1}{N}\frac{dN}{dt} = \frac{1}{E}\frac{dE}{dt} \tag{S9}$$

Equivalently, we can rewrite this in terms of the quotas $Q_N$ and $Q_E$.

$$\frac{dQ_N}{dt} = \frac{dQ_E}{dt} = 0 \tag{S10}$$

We solve equation A10 to get two sets of steady-state quota values. The first set corresponds to a dead population where $\tilde{Q}_{E,1} = 0$ and $\tilde{Q}_{N,1} = \frac{Q_{str}}{1-f_{nut}}$. The second set of non-trivial $\tilde{Q}_N$ and $\tilde{Q}_E$ is:

$$\tilde{Q}_{N,2} = \frac{e_\omega^2\, f_{\mathrm{PPP}}^2\, P_0\, Q_{\mathrm{str}} + f_{\mathrm{nut}}\,(1 - f_{\mathrm{PPP}})\, V_0\, \phi_\rho}{e_\omega\, \phi_N\, f_{\mathrm{PPP}}\,(e_\omega\,(1 - f_{\mathrm{nut}}) f_{\mathrm{PPP}}\, P_0 + f_{\mathrm{nut}}\, V_0\, \phi_N)} \tag{S11}$$

$$\tilde{Q}_{E,2} = \frac{P_0\,\left((1 - f_{\mathrm{nut}} - f_{\mathrm{PPP}} + f_{\mathrm{nut}}\, f_{\mathrm{PPP}})\phi_\rho - e_\omega\, f_{\mathrm{PPP}}\, Q_{\mathrm{str}}\, \phi_N\right)}{e_\omega\,(1 - f_{\mathrm{nut}}) f_{\mathrm{PPP}}\, P_0 + f_{\mathrm{nut}}\, V_0} \tag{S12}$$

Using the parameters in Appendix A5, we plot $\tilde{Q}_N$ and $\tilde{Q}_E$ against $f_{\mathrm{PPP}}$ and $f_{\mathrm{nut}}$ in Fig.2 in the main text. The top right corner of both plots in Fig. 2 corresponds to the trivial steady-state solution ($\tilde{Q}_{N,1}, \tilde{Q}_{E,1}$) where the population is extinct. Going forward, we focus on the non-trivial solution and refer to it as $\tilde{Q}_N = \tilde{Q}_{N,2}$ and $\tilde{Q}_E = \tilde{Q}_{E,2}$ for brevity.

We substitute $Q_N = \tilde{Q}_N$ into eq. A5 to calculate exponential growth rate $\tilde{\mu}$ plotted in Fig. 3 in the main text. For Fig. 3a, we use the FindMaximum command of Wolfram Mathematica to numerically calculate ($\tilde{f}_{\mathrm{PPP}}^{\mathrm{opt}}, \tilde{f}_{\mathrm{nut}}^{\mathrm{opt}}$), the allocations that maximize the exponential growth rate. This point is marked in black in Fig. 3a.

We also calculate ($\tilde{f}_{\mathrm{PPP}}^{\mathrm{opt}}, \tilde{f}_{\mathrm{nut}}^{\mathrm{opt}}$) analytically by calculating the critical points of growth, values of $f_{PPP}$ and $f_{nut}$ where the partial derivatives of the steady-state growth rate $\tilde{\mu}$ w.r.t $f_{PPP}$ and $f_{nut}$ are zero ($\nabla\tilde{\mu}(f_{PPP}, f_{nut}) = 0$). Since $f_{PPP}$ and $f_{nut}$ are fractions, the critical points must also satisfy $(f_{PPP}, f_{nut}) \in [0,1]$. Finally, we verify that the critical point is a local maximum by checking that the determinant of the Hessian matrix of the exponential growth function ($\mu$) is positive ($|\mathbf{H}_\mu| > 0$) and that the top left element of the Hessian is negative ($(\mathbf{H}_\mu)_{1,1} < 0$). The complete calculations can be seen in the accompanying Mathematica notebook.

#### 2. Modulation of optimal allocations in response to changes in external nitrogen and light environments

Here, we examine how the optimal allocations $(\tilde{f}_{\text{PPP}}^{\text{opt}}, \tilde{f}_{\text{nut}}^{\text{opt}})$ depend on changing external environments. We start with analyzing $\left(\tilde{f}_{\text{PPP}}^{\text{opt}}, \tilde{f}_{\text{nut}}^{\text{opt}}\right)$as a function of the potential uptake rate ratio (Fig. A1a). We divide both the numerator and denominator of the steady-state quota expressions $(\tilde{Q}_N, \tilde{Q}_E)$ by $V_0$ and collect all $P_0$ and $V_0$ terms as $P_0/V_0$, thus acquiring the steady-state quota expressions in terms of $P_0/V_0$. We then substitute the quota expressions in terms of $P_0/V_0$ and re-calculate the critical points of the steady-state growth function $(\nabla\tilde{\mu}(f_{\text{PPP}}, f_{\text{nut}}) = 0)$. Imposing the constraint $(f_{\text{PPP}}, f_{\text{nut}}) \in [0,1]$ on the critical points gives us the optimal allocations $\left(\tilde{f}_{\text{PPP}}^{\text{opt}}, \tilde{f}_{\text{nut}}^{\text{opt}}\right)$ in terms of $P_0/V_0$. The complete calculations can be seen in the accompanying Wolfram Mathematica notebook.

To maintain balanced growth, the cell adjusts its optimal allocations to account for mismatches in the carbon vs nitrogen uptake rates. An increase in the ratio of potential uptake rates of carbon and nitrogen leads to an increase in the optimal $f_{\text{nut}}$ and a decrease in the optimal $f_{\text{PPP}}$. When $P_0/V_0$ is low (left side of Fig. A1a), the carbon uptake is small compared to the nitrogen uptake. Therefore, the cell allocates a high amount of carbon to $f_{\text{PPP}}$ (blue curve) to maximise carbon uptake and minimise carbon loss via respiration. The optimal nitrogen allocation to uptake ($f_{\text{nut}}$, yellow curve) is low so that the fraction of nitrogen allocated to photosynthesis ($f_{\text{photo}}$) and consequently carbon uptake is maximised. It is instructive to consider the limiting case of $\frac{P_0}{V_0} \to 0$, where the nitrogen intake rate is infinitely high ($V_0 \to \infty$). In response, the optimal allocation of nitrogen to uptake tends towards 0 ($f_{\text{nut}} \to 0$). The optimal allocation of carbon towards PPP also tends towards a maximum value. The cell always needs some carbon for respiration to produce energy, so $f_{\text{PPP}}$ cannot be 1 even in this limiting case.

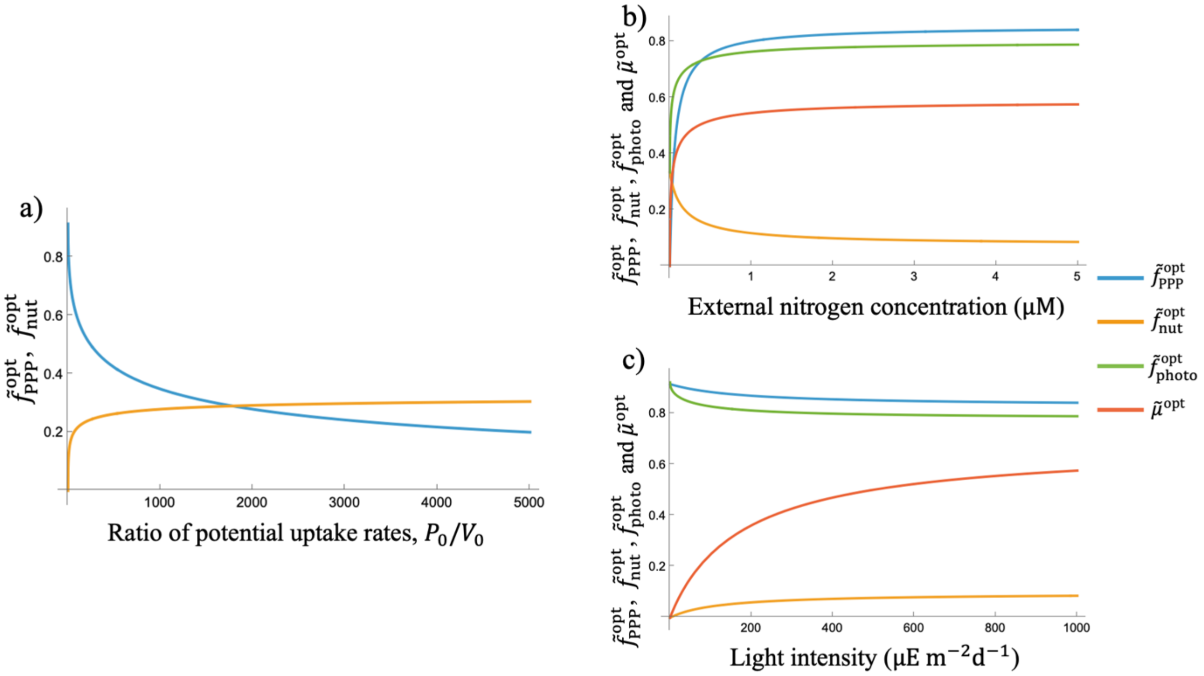


*Figure S1: a) Optimal allocations of carbon to PPP ($\tilde{f}_{PPP}^{opt}$, blue curve) and nitrogen to uptake ($\tilde{f}_{nut}^{opt}$, yellow curve) as a function of the ratio of potential uptake rates ($P_0/V_0$) of carbon ($P_0$) and nitrogen ($V_0$). b) Light-saturated, nutrient-limited growth: $\tilde{f}_{PPP}^{opt}$ (blue curve), $\tilde{f}_{nut}^{opt}$ (yellow curve), $\tilde{f}_{photo}^{opt}$ (green curve) and growth rate ($\tilde{\mu}^{opt}$, red curve) as a function of external nitrogen concentration. c) Light-limited, nutrient-saturated growth: $\tilde{f}_{PPP}^{opt}$ (blue curve), $\tilde{f}_{nut}^{opt}$ (yellow curve), $\tilde{f}_{photo}^{opt}$ (green curve) and growth rate ($\tilde{\mu}^{opt}$, red curve) as a function of irradiance. Note that the loss rate $l_C = 0$ for these plots.*

At high $P_0/V_0$ values (right side of Fig. S1a), the carbon uptake rate is large compared to the nitrogen uptake. Therefore, the optimal allocation of C to PPP ($f_{PPP}$, blue curve) decreases leading to high respiration, high energy generation and consequently higher levels of nitrogen uptake. The optimal nitrogen allocation to uptake $f_{nut}$ (yellow curve) also increases to maximize nitrogen intake. Note that the rate of increase of optimal $f_{nut}$ slows down with increasing $P_0/V_0$ since as well. As earlier, we consider the limiting case of $\frac{P_0}{V_0} \to \infty$ where the carbon intake rate $P_0 \to \infty$. To counter the infinitely high carbon intake, the carbon allocation to PPP, $f_{PPP}$ approaches zero ($f_{PPP} \to 0$). Since the cell always needs some nitrogen for photosynthesis, optimal $f_{nut}$ tends towards a maximum value that is lower than 1.

Under both extreme scenarios ($P_0 \to \infty$ and $V_0 \to \infty$), the high uptake rates are counteracted by setting the corresponding allocation ($f_{PPP}$ for $P_0 \to \infty$ and $f_{nut}$ for $V_0 \to \infty$) to extremely low levels. However, the upper bound for both allocation parameters is always less than 1, reflecting the importance of keeping respiration and photosynthesis going even under extreme circumstances.

Next, we relax the assumptions of light and nutrient saturation and calculate optimal steady-state growth dependent on external light intensity and nutrient concentration. To do this, we simply replace the potential uptake rate of carbon ($P_0$) with light-dependent uptake $P$ in the expression for optimal allocations $\left(\tilde{f}_{\text{PPP}}^{\text{opt}}, \tilde{f}_{\text{nut}}^{\text{opt}}\right)$:

$$P = P_0\, f_{\text{photo}} \frac{I}{K_I + I} \tag{A13}$$

Here, $I$ denotes the light level and $K_I$ is the half-saturation constant. We currently assume no photo-inhibition, however incorporating photoinhibition would be straightforward in the model. We also replace the potential uptake rate of nitrogen ($V_0$) with nutrient-dependent uptake $V$ in the expression for optimal allocations $\left(\tilde{f}_{\text{PPP}}^{\text{opt}}, \tilde{f}_{\text{nut}}^{\text{opt}}\right)$:

$$V = V_{\max}[f_{\text{nut}}, Q_E] \frac{S}{K_S + S} \tag{A14}$$

As a result, we get the optimal allocations $\left(\tilde{f}_{\text{PPP}}^{\text{opt}}, \tilde{f}_{\text{nut}}^{\text{opt}}\right)$ in terms of the external nutrient concentration $S$ and light level $I$. Using the optimal allocations $\left(\tilde{f}_{\text{PPP}}^{\text{opt}}, \tilde{f}_{\text{nut}}^{\text{opt}}\right)$, we also calculate the resulting allocation to $f_{\text{photo}}$ and the consequential optimal steady-state growth $\tilde{\mu}^{\text{opt}}$. The complete expression for the optimal steady-state growth rate is provided in the accompanying Mathematica notebook. We explore the results of these analyses by looking at them in order of increasing complexity: only nutrient-limited growth ($I = 1000$), only light-limited growth ($S = 5$) and both nutrient- and light-limited growth.

*Light-saturated and nutrient-limited growth*

In Fig. S1b, we plot the optimal allocation parameters $f_{\text{PPP}}$ (blue curve) and $f_{\text{nut}}$ (yellow curve) as a function of changing external nitrogen concentrations while light levels are held constant at a saturating level ($I = 1000$). Moving from left to right in Fig. S1b corresponds to moving from right to left in Fig. A1a. As external nutrient concentrations increase, nutrient uptake increases leading to a saturating decline in nitrogen allocation to uptake $f_{\text{nut}}$ (yellow curve in Fig. S1b) and a corresponding saturating increase in nitrogen allocation to photosynthesis ($f_{\text{photo}}$, green curve in Fig. S1b). The carbon allocation to PPP ($f_{\text{PPP}}$, blue curve in Fig. S1b) also increases to maximize carbon assimilation and minimize respiratory carbon loss. Growth ($\tilde{\mu}^*$, red curve in Fig. S1b) is a product of photosynthesis rate ($P = P_0\, f_{\text{photo}}$) and the carbon allocation to PPP $f_{\text{PPP}}$, so it increases in a saturating manner with increasing external nitrogen concentration. Since growth is a product of two saturating functions, it saturates quickly. Note that external nitrogen concentration determines internal allocation of carbon and nitrogen that then determines growth.

*Light-limited and nutrient-saturated growth*

We now consider the converse situation where external nitrogen concentration is saturating ($S = 5$), and growth is light-limited. We plot the optimal values of $f_{\text{PPP}}$ and $f_{\text{nut}}$ against increasing light levels in Fig. S1c. Increasing light levels by moving from left to right in Fig. S1c is equivalent to moving from left to right in Fig. A1a. As light levels increase, carbon uptake rates go up and the optimal allocation to PPP, $f_{PPP}$ (blue curve in Fig. S1c) goes down in a saturating manner. This also leads to an increase in respiration and energy production which then leads to an increase in nitrogen uptake rate. The nitrogen allocation to uptake $f_{\text{nut}}$ (yellow curve in Fig. S1c) also increases in a saturating manner with external light levels, consequently leading to a saturating decline in nitrogen allocation to photosynthesis ($f_{\text{photo}}$ red curve in Fig. S1c).

Growth ($\tilde{\mu}^{\text{opt}}$) has three components: the externally driven carbon uptake $\left(P_0 \frac{I}{K_I+I}\right)$, the nitrogen allocation to photosynthesis ($f_{\text{photo}}$) and the carbon allocation to PPP ($f_{\text{PPP}}$). While both $f_{\text{photo}}$ and $f_{\text{PPP}}$ decline in a saturating fashion with increasing light levels, the externally driven carbon uptake $\left(P_0 \frac{I}{K_I+I}\right)$ increases in a saturating fashion and drives the saturating increase of growth (red curve in Fig. S1c) with light. In contrast to nitrogen, light impacts growth both directly through carbon uptake $\left(P_0 \frac{I}{K_I+I}\right)$ and indirectly through internal allocation to $f_{\text{photo}}$ and $f_{\text{PPP}}$. The direct impact of light dominates the growth response, therefore light-limited growth does not saturate as quickly as nutrient-limited growth.

#### 3. Growth vs nitrogen and energy quotas

Next, we analyze the impact of internal storage of nutrients and energy on growth. To do this, we go back to analyzing the model under a nutrient- and light-saturated environment. In Section 2, we calculated both optimal growth ($\tilde{\mu}^{\text{opt}}$) and the steady-state quotas ($\tilde{Q}_N, \tilde{Q}_E$) in terms of the ratio of potential uptake rates $P_0/V_0$. Here, we vary $P_0/V_0$ as a common parameter and plot optimal growth ($\tilde{\mu}^{\text{opt}}$) against nitrogen quota ($\tilde{Q}_N$, Fig. S2a) and energy quota ($\tilde{Q}_E$, Fig. S2b) in two parametric plots. Since $\tilde{Q}_N$ and $\tilde{Q}_E$ are linked through balanced growth, not all combinations of $\tilde{Q}_N$ and $\tilde{Q}_E$ are feasible at steady-state. Plotting growth parametrically through varying $P_0/V_0$ ensures that only feasible $\tilde{Q}_N$ and $\tilde{Q}_E$ values are used for plotting.

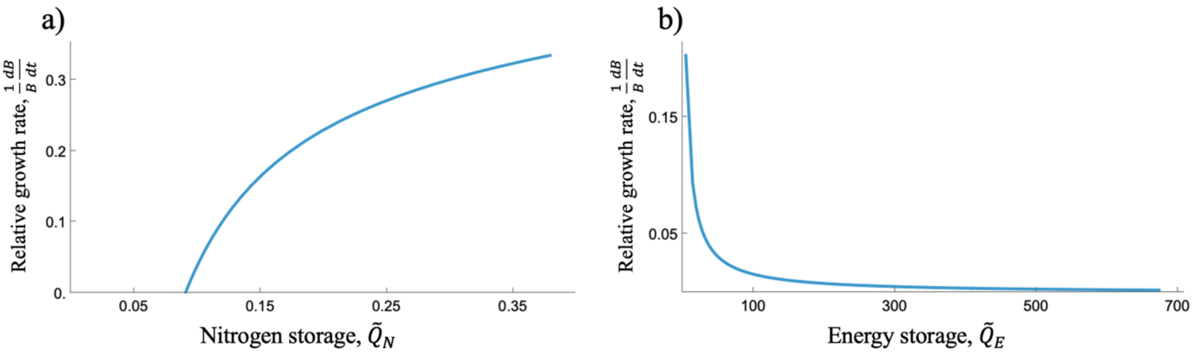


*Figure S2: Exponential growth rate ($\frac{1}{B}\frac{dB}{dt}$) as a function of internal storage of a) nitrogen, $\tilde{Q}_N$ and b) energy, $\tilde{Q}_E$.*

In Fig. S2a, we plot the common log of optimal exponential growth rate (y-axis) as a function of nitrogen storage $\tilde{Q}_N$ (x-axis). Exponential growth rate increases with nitrogen storage in a Droop-like manner (Fig. S2a), with a minimum $Q_N$ value needed for growth. Using a simpler model where only nitrogen was stored in the cell, Pahlow and Oschlies (2013) showed that the Droop relationship could be derived from optimal allocation of nitrogen. In contrast, our model considers the storage of both nitrogen and energy. This restricts feasible $Q_N$ values at steady-state in our model. Further, our model considers optimal allocations of both carbon and nitrogen. Despite the added complexity, we are able to recover a Droop-like shape for growth vs nitrogen quota.

In Fig. S2b, we plot the exponential growth rate as a function of energy storage $\tilde{Q}_E$. The exponential growth rate decreases hyperbolically as energy storage $\widetilde{Q_E}$ increases. Energy is produced by respiration, so low energy levels imply that most carbon is directed to PPP thus resulting in high levels of growth. Low energy quota values also imply high levels of consumption by nutrient uptake. So, low energy quota corresponds with high levels of $\tilde{Q}_N$ (right side of Fig. S2a) that leads to a saturation in growth. Note that while energy quotas are low, energy quota never hits zero since the cell always needs to direct some carbon through respiration to generate energy for nutrient uptake. High energy quota results from a high allocation of carbon to respiration, thus reducing growth rates. High energy quotas also imply low energy consumption through nutrient uptake and therefore correspond to the lower end of $\tilde{Q}_N$ values in Fig. S2a.

#### 4. Model analysis in a chemostat

Next, we analyze the model dynamics under equilibrium in a chemostat where the phytoplankton growth rate is controlled by the dilution rate, $D$. Phytoplankton take up nitrogen ($S$) following a Monod-type response from eq. S14. The chemostat has a supply rate $S_{in}$. The nitrogen dynamics can be written as:

$$\frac{dS}{dt} = D\,(S_{\text{in}} - S) - V_{\max}[f_{\text{nut}}, Q_E]\frac{S}{K_S + S}\,C \qquad \text{(S15)}$$

In a chemostat, the model can thus be written as a combination of external nitrogen dynamics (eq. S15) along with the internal carbon, nitrogen and energy dynamics (eq. S1, S17 and S8). We set all loss rates equal to the dilution rate: $l_C = l_N = l_E = D$. We set the time derivatives to zero and solve for equilibrium values. We find two trivial solutions where the population goes extinct and a non-trivial solution. In the non-trivial solution, we get:

$$Q_N^* = \frac{e_\omega f_{\text{PPP}} P_0 Q_{\text{str}}}{e_\omega f_{\text{PPP}} P_0 (1 - f_{\text{nut}}) - D}, Q_E^* = \frac{1 - f_{\text{PPP}}}{e_\omega f_{\text{PPP}}}\phi_\rho - \phi_N Q_N^*, S^* = \frac{K_S V^*}{\frac{V_0}{\phi_N} - V^*}, C^* = \frac{S_{in} - S^*}{Q_N^*} \qquad \text{(A16)}$$

Here, $V^* = \frac{D\,Q_N^*}{f_{\text{nut}} Q_E^*}$ can be interpreted as the equilibrium nitrogen uptake rate. Fig. 5 in the main text explores how $Q_N^*$, $Q_E^*$ and $S^*$ vary as a function of the allocation parameters

$f_{\text{PPP}}$ and $f_{\text{nut}}$. For Fig. 5c, we use the FindMinimum command in Wolfram Mathematica to numerically calculate $(f_{\text{PPP}}^{*,\text{opt}}, f_{\text{nut}}^{*,\text{opt}})$ (red point in Fig. 5c), the optimal allocation strategy that minimizes $S^*$.

We also analytically calculate the optimal competitive strategy $(f_{\text{PPP}}^{*,\text{opt}}, f_{\text{nut}}^{*,\text{opt}})$. First, we solve $\nabla S^*(f_{\text{PPP}}, f_{\text{nut}}) = 0$ to obtain critical points. We get five solutions but imposing the constraint of $(f_{PPP}, f_{nut}) \in [0,1]$ results in only one biologically feasible critical point:

$$f_{\text{PPP}}^{*,\text{opt}} = \frac{(5D + \Psi)\phi_\rho - \Gamma}{2\,\Delta}, f_{\text{nut}}^{*,\text{opt}} = \frac{2\,\phi_\rho(\Psi - D) - \Gamma}{3\,\Psi\,\phi_\rho} \quad (S17)$$

Here, $\Psi$, $\Gamma$ and $\Delta$ are three intermediate variables defined as:

$$\Psi = e_\omega\, P_0, \Theta = e_\omega^2\, P_0 \text{Q}_{\text{str}} \Phi_{\text{N}},$$
$$\Delta = (2D + \Psi)\phi_\rho - \Theta, \Gamma = \sqrt{\phi_\rho^2 (D - \Psi)^2 + 12\, D\, \Theta\, \phi_\rho} \quad (S18)$$

Using the parameter values from Table S1 below, we numerically verify that the critical point is a minimum for a range of $D$ where the equilibrium is feasible. We then plot $(f_{\text{PPP}}^{*,\text{opt}}, f_{\text{nut}}^{*,\text{opt}})$ against the dilution rate $D$ in Fig. 6 in the main text to understand the effect of dilution rate on the optimal allocations. All calculations were done in Wolfram Mathematica and can be seen in the accompanying Mathematica notebook here: https://github.com/raviranjan545/Carbon_Budget_Model.

*Table S1: Parameter values used in the main text.*

| Parameter | Value | Units | Definition |
|---|---|---|---|
| $Q_{str}$ | 0.03 | $\text{mol N (mol C)}^{-1}$ | Nitrogen quota allocated to structure |
| $K_S$ | 1 | $\mu\text{M}$ | Half-saturation constant of nutrient uptake |
| $V_0$ | 1.2 | $\text{day}^{-1}$ | Potential maximum nitrogen uptake rate |
| $S_{in}$ | 10 | $\mu\text{M}$ | Nitrogen inflow concentration; used in chemostat analysis |
| $a$ | 0.2 | $\text{day}^{-1}$ | Dilution rate; used in chemostat analysis |
| $K_I$ | 200 | $\mu\text{E m}^{-2}\text{ s}^{-1}$ | Half-saturation constant of photosynthesis |
| $P_0$ | 2.6 | $\text{day}^{-1}$ | Potential photosynthesis rate |
| $e_\omega$ | 0.4 | unitless | Efficiency of carbon assimilation of the PPP |
| $\phi_N$ | 3 | $\text{mol ATP (mol N)}^{-1}$ | mol energy (ATP) used per mol of N taken up |

| | | | |
|---|---|---|---|
| $\phi_\rho$ | 5 | $\text{mol ATP (mol C)}^{-1}$ | mol energy (ATP) generated per mol of C respired |
| $l_C$ | 0.2 | $\text{day}^{-1}$ | Carbon loss rate |
| $l_N$ | 0.2 | $\text{day}^{-1}$ | Nitrogen loss rate |
| $l_E$ | 0.2 | $\text{day}^{-1}$ | Energy loss rate |
